# Fishing for biodiversity by balanced harvesting

**DOI:** 10.1101/2021.06.27.450047

**Authors:** Richard Law, Michael J Plank

## Abstract

Fisheries are damaging, and seemingly incompatible with the conservation of marine ecosystems. Yet fish are an important source of food, and support the lives of many people in coastal communities. This paper considers an idea that a moderate intensity of fishing, appropriately scaled across species, could help in maintaining biodiversity, rather than reducing it. The scaling comes from an intuition that rates of fishing mortality of species should be kept in line with production rates of the species, a notion known as balanced harvesting. This places species conservation and exploitation on an equal footing in a single equation, showing quantitatively the relative levels of fishing mortality that species of different abundance can support. Using a dynamic model of a marine ecosystem, we give numerical evidence showing for the first time that fishing, if scaled in this way, can protect rarer species, while allowing some exploitation of species with greater production. This works because fishing mortality rates, when scaled by production, are density-dependent. Such fishing, operating adaptively to follow species’ production rates over time, contains a feedback that would help to protect species from overfishing in the presence of uncertainty about how marine ecosystems work.

## 1 Introduction

Conservation and fisheries coexist uneasily (Salomon et al., 2011; Garcia et al., 2014). Fisheries have many destructive effects, and make a strong case for reserves to protect marine ecosystems from fishing (Lubchenco and Grorud-Colvert, 2015). The motivation for this paper is not to question the conservation value of marine protected areas, but rather to consider how fishing could be made more compatible with the goals of conservation, where complete closure of fisheries is not an option. This is important because there is potential for serious conflict between the stakeholders of conservation and those of fisheries (Garcia et al., 2003). On one hand, the lives and livelihoods of many people in coastal areas of the world depend on food from the sea. On the other hand, we are the custodians of marine ecosystems, and have a responsibility to leave them undamaged for the future. Schemes that allow moderate exploitation, while assisting the conservation of marine ecosystems, are therefore worth considering.

Such fishing schemes need to be organised at the ecosystem level, and the literature contains various ideas about fishing at this broad scale. For instance, Larkin (1977) suggested that maximum sustainable yields (MSYs) could be applied to aggregated assemblages of fish species rather than to single species, harvesting all species above a common body size. Variations on the theme of multispecies MSY have subsequently been developed including, for example, ‘a pretty good multispecies yield’ (Rindorf et al., 2017). Fowler (1999) argued that natural predators of exploited fish stocks could indicate the sustainable ways in which to apply fishing mortality; see also Caddy and Sharp (1986). Kleisner and Pauly (2011) proposed ‘fishing in balance’ to deal with fisheries-induced trophic changes in ecosystems (fishing down the food web: Pauly et al., 1998). Rehren and Gascuel (2020) went further, proposing ‘balanced structure harvesting’, to minimise the disruption to trophic structure.

Trophic structure, while important, is one of a number of ways in which marine ecosystems are impacted by fishing. Biodiversity is another, as is size structure within species. Garcia et al. (2012) suggested ‘balanced harvesting’ (BH) as a guiding principle for exploitation at this more nuanced level. The idea of BH is to bring fishing mortality more in line with natural productivity of species and body sizes (reviewed by Heath et al., 2017; Zhou et al., 2019). This builds on an intuition that the death rate from fishing should correspond to the rate at which species and body sizes can replace the biomass extracted by fishing. Fishing in this way should help protect ecosystem components that are rare (low biomass), and those that have low somatic growth rates.

BH is an ecosystem approach to fishing. Its organising principle, productivity, depends on fish growth, which comes from fish feeding on one another and on plankton (causing prey death). Such feeding depends first on the size of prey items relative to the consumer, and secondarily on their taxonomic identities (Jennings et al., 2001), thereby coupling species together in a manner that depends on their body-size distributions (Persson et al., 2014). There are some tools for studying ecosystem dynamics at this fine level. These include multispecies, size-spectrum models which track body-size distributions of species as they change over time, one size distribution for each species (Andersen and Beyer, 2006; Hartvig et al., 2011; Blanchard et al., 2014), and we use such models here. Importantly they have, at their heart, the core ecosystem coupling, that fish growth and, to a large extent death, follows directly from predators feeding on prey. This means that body growth is determined dynamically within the ecosystem, eliminating the need for an external assumption about growth, such as a von Bertalanffy equation. From this coupling, the flow of biomass through components of the ecosystem emerges in a quantifiable way (Law et al., 2016).

Widely in marine biology, ‘productivity’ is taken to be a gain in mass per unit area per unit time (dimensions: M L^−2^ T^−1^); the term ‘primary productivity’ is an instance of this. However, in the BH literature there are currently two different meanings of the term productivity, one being a total rate of production as above (M L^−2^ T^−1^), and the other being a rate per unit biomass (dimensions: T^−1^) (Heath et al., 2017; Zhou et al., 2019; Nilsen et al., 2020). The two meanings have entirely different consequences in the context of BH. This is because the former sets the fishing mortality rate *F* in proportion to a production rate *P* (*F* ∝ *P*), whereas the latter sets *F* ∝ *P/B* where *B* is biomass. These two versions of BH need to be treated separately. To avoid confusion in this paper, we distinguish between them using indices BH_*P*_, BH_*P/B*_, and refer to *P* as production rate.

The results in this paper show for the first time that BH can have clear benefits for the maintenance of biodiversity. These benefits come from setting *F* ∝ *P* (BH_*P*_). They do not come from setting *F* ∝ *P/B* (BH_*P/B*_). BH_*P*_ thus provides a way to reconcile some objectives of conservation with some of fisheries, and suggests a way in which moderate exploitation of marine ecosystems could be brought better in line with the needs of conservation. The distinction between BH_*P*_ and BH_*P/B*_ is important in practice, because traditional fisheries management leans towards BH_*P/B*_, for instance in the assumption *F* = *M* of the length-based proxy for *F*_*MSY*_ (ICES, 2021, Annex 3).

To get these results, we used dynamic models that hold in place the general empirical relationship between *B* and *P* in marine ecosystems. The modelling approach is justified by the fact that the basic features of BH should be generic and independent of the exact choice of ecosystem. Moreover, in real-world marine ecosystems, data collection is often focused on the major commercial fish stocks, and not on rarer non-commercial species, some of which may be vulnerable to decisions made about fisheries management. The aim of this work was to put conservation and exploitation on an equal footing, and to investigate management strategies that could potentially deliver benefits for both conservation and fisheries. We therefore needed to give all species the same attention, whether common or rare, and modelling is the best tool for doing so. Model conclusions of course still need to be tested and refined over time in empirical settings.

## 2 Methods

We explain here how we implemented a model of multispecies ecosystem dynamics and the harvesting patterns applied to it. The results can be understood without going into details of the model, but it is important to know they are underpinned by a rigorous mathematical framework. The model was designed to contain a range of taxa from microscopic photosynthetic plankton (primary producers) to large fish species, coupled throughout by feeding, and calibrated to hold in place some basic properties of biomass and production rate observed in marine ecosystems.

### 2.1 Life histories

We used a common template for fish life histories (Fig. 1). In this template, fish of all species start life as eggs of mass 1 mg, and grow as a result of feeding on smaller organisms, process (a) in Fig. 1, as defined by a feeding kernel. Thus, in their early life stages, the fish feed on plankton. As they grow larger, their diet shifts gradually from plankton towards smaller fish of their own and other species. These fish are themselves at risk of being eaten by larger fish (process b). In addition to death from predation, we include a risk of death from other causes (process c) which is especially high in the larval stage, and death from fishing (process d). Reproduction (process e), starts when fish reach a body mass around 10% of their maximum body mass, and entails an increasing transfer of incoming food to egg production, as opposed to further somatic growth. The maximum body mass for a given species is reached when all incoming food is allocated to egg production and none to somatic growth. From this template, arbitrary life histories for fish species were created by assigning a random values to the maximum body mass, together with some modest random variation between species in larval death rates. The template could be modified as required to represent the life histories of multicellular taxa other than teleost fish.

**Figure 1:**
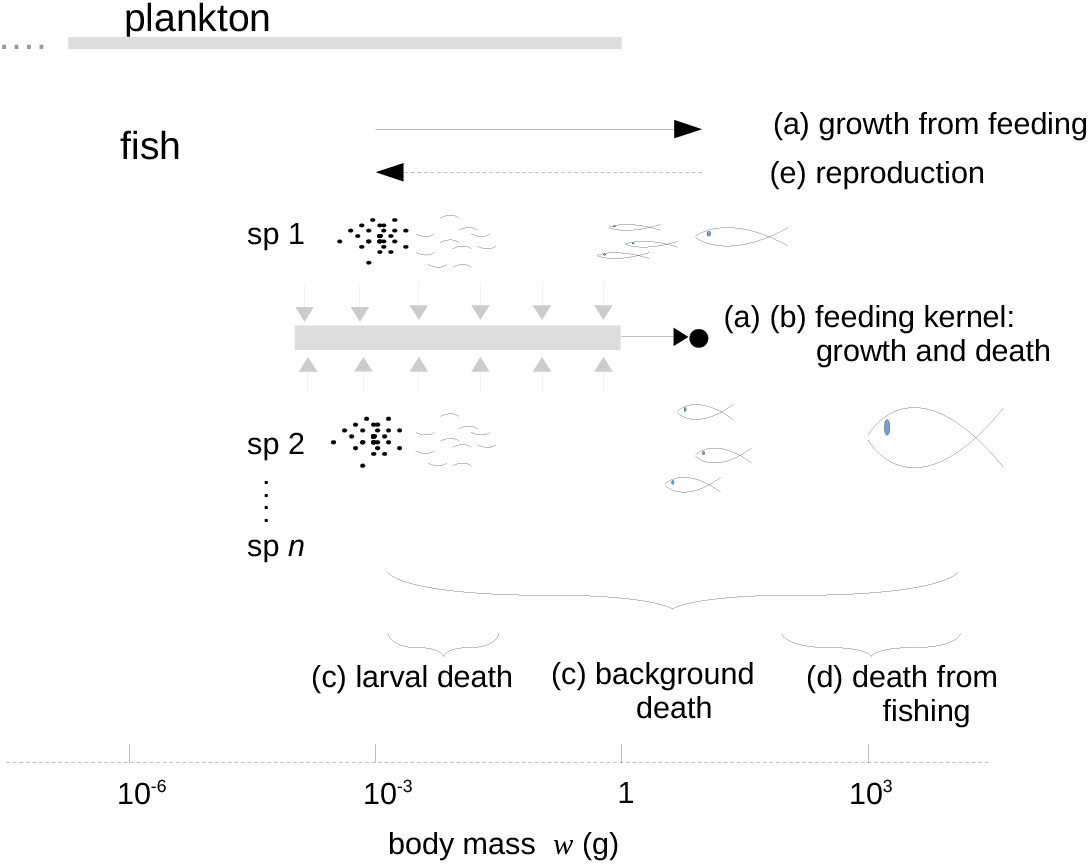
Template from which a set of *n* fish species with a range of life histories was constructed by varying maximum body mass *w*_∞_ and larval death rate. The feeding kernel sets a body mass range of prey items eaten relative to the body mass of the consumer, and is sketched as the grey bar for a fish at the body mass shown by the black circle. The kernel is tethered to the consumer, and ‘moves’ to the right as a result of the growth that follows from eating this food.

### 2.2 Size-spectrum dynamics

The state variables used in the ecosystem model are densities of individuals. The densities of individuals of type *i* and body mass *w* are denoted *ϕ*_*i*_(*w, t*) at time *t*. The fish assemblage is disaggregated to an arbitrary number *n* of species indexed *i* = 1, …, *n*. However, the potentially complicated plankton assemblage is amalgamated into a single density function indexed 0, and is described separately below.

The basic processes sketched in Fig. 1 are given in a model of multispecies size-spectrum dynamics as

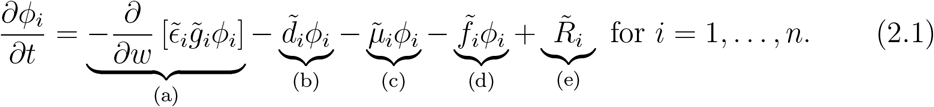

Eq. (2.1) is a multi-species, size-structured form of the McKendrick–von Foerster equation (McKendrick, 1926; von Foerster, 1959; Silvert and Platt, 1978). This equation tracks the evolution of the size structure and biomass of each species over time. The terms in Eq. (2.1) represent: (a) somatic growth; (b) predation mortality; (c) other natural mortality; (d) fishing mortality; and (e) reproduction. A detailed definition of these terms is provided in Appendix A, here we give an informal description.

Eq. (2.1) is derived from a book-keeping of biomass as it moves from one species and size class to another as a result of processes (a)–(e). This means that, as in real marine ecosystems, species are coupled together through their feeding. Growth in body mass or reproduction of a consumer is necessarily accompanied by death of its prey. The derivations of the mass-specific food intake rate 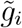, reproduction rate 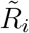, and predation mortality rate 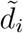 given in Appendix A reflect this fundamental coupling: 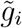 and 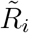 are functions of the abundance of prey of suitable size and species; 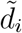is a function of the abundance of predators of suitable size and species. The function 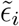 partitions the food between somatic growth and reproduction according to species and body size. In this way fish growth is internalised in the dynamics, and is independent of any external model such as a von Bertalanffy growth equation.

In addition to these internal biomass flows, biomass enters the system through primary production by plankton, and leaves the system as a result of inefficient feeding, a non-predation natural mortality rate 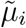 and a fishing rate 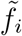. Nonpredation natural mortality is assumed to be composed of a general background mortality rate which includes senescence, and an additional mortality term in the larval stage. This extra term is not usually included in size-spectrum models, but there is an important feedback between larval mortality and body growth which needs to be made explicit (Ricker and Foerster, 1948; MacCall, 1980; Canales et al., 2020). The feedback works as follows. When food is plentiful, growth through the larval stage is fast and larval mortality is relatively small. When food becomes scarce, growth through the larval stage slows down, and larval mortality becomes relatively large.

Plankton were taken to span a mass range from 10^−1^0 g to 1 g, the function *ϕ*_0_(*w, t*) being the density at size *w* and time *t*. Over most of the size range, plankton are unicellular, meaning that a somatic growth process like that in Eq. (2.1) is not needed. We therefore modelled the dynamics of *ϕ*_0_(*w, t*) via a simpler logistic dynamic

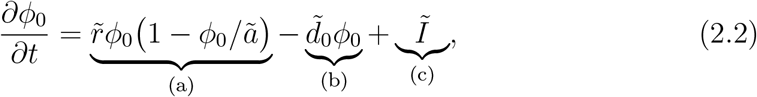

where term (a) contains the intrinsic rate of increase 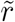 and carrying capacity *ã* of the logistic model, term (b) contains the mortality rate 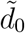 due to predation by fish, and term (c) is an immigration rate *I* independent of local dynamics. These rates are all body-size dependent. (Details of the terms are in Appendix A.)

Together, Eqs (2.1) and (2.2) describe the dynamics of an exploited, multispecies, marine ecosystem. The equations disaggregate the ecosystem down to fish species and then down to body mass within fish species, and are built on the book-keeping of biomass as it flows from primary producers up through the food chain. This means that, as in real marine ecosystems, species are coupled together through their feeding. Once parameter values have been specified (Appendix C), the dynamics can be simulated numerically.

### 2.3 Patterns of exploitation

BH sets fishing mortality rates in accordance with production rate *P* (dimensions: M L^−2^ T^−1^) and biomass *B* (dimensions: M L^−2^), which are species-level properties of longstanding interest in fisheries science (Allen, 1971; Dickie, 1972). In size-spectrum models, production emerges from body growth of consumers eating prey, and we therefore measured it directly from the dynamic variables which describe this biomass flow (see below). This is in contrast to logistic-like biomass models in which body growth is either absent, or specified by a life history which is independent of the prey needed for somatic growth to take place (de Kerckhove, 2015; Zhou and Smith, 2017; Plank, 2018). It is also in contrast to equilibrium models which use an equality *P* = *ZB* (where *Z* is the total rate of mortality) on the grounds that, at equilibrium, the gain in biomass from production must balance the loss of biomass through mortality (Dickie, 1972). The direct measure of biomass flow needs some care, as it has to deal with: (1) the body-mass range over which *B* and *P* are measured; (2) the partition of incoming mass between somatic growth and reproduction; (3) effects on *P* of biomass flow into and out of the measured body-size range; (4) body-size-dependent variations in growth rate and death rate (natural and fishing). Note that measurements of *P* and *B*, when constructed directly from biomass flows described by Eq. (2.1), apply to ecosystems which are not at equilibrium, as well as to those which are at equilibrium.

The biomass per unit area of species *i* at time *t* in a small range of body mass [*w, w* + *dw*], was written as *b*_*i*_(*w, t*)*dw* = *wϕ*_*i*_(*w, t*)*dw* (dimensions: M L^−2^). The total biomass per unit area in a range of body mass bounded below and above by *<u>w</u>*_*i*_ and 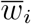 respectively is then

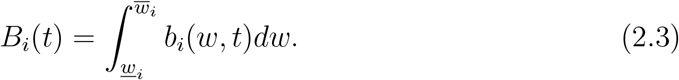

Similarly, the somatic production rate *P*_*i*_(*t*) per unit area per unit time over the same size range was taken as:

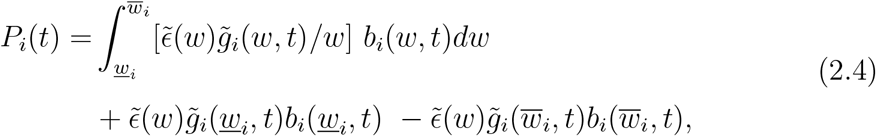

written in this way to emphasise the fact that production rate depends on biomass. The boundary terms at 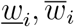 deal with the flow of biomass into and out of the size range over which *P*_*i*_ is measured. If the lower bound is at the size of eggs, the flow-in term becomes the rate at which egg mass is produced, which is equal to 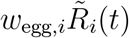.

Yield from fishing had a similar general form

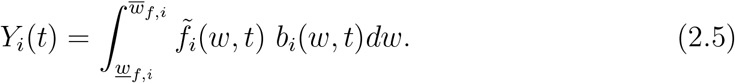

where 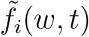 is the rate of fishing mortality on species *i* at body mass *w* and time *t*, as in Eq. (2.1), with minimum and maximum sizes as 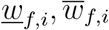 respectively for species *i*. For simplicity we assumed that all fish entered a mixed-species fishery at a single size *<u>w</u>*_*f*_ and all fish of species *i* with body mass greater than *<u>w</u>*_*f*_ were caught at the same rate *F*_*i*_(*t*) from this size onwards (the rate potentially being species- and time-dependent *t*.) Under these conditions, fishing mortality can be factored out of the integral

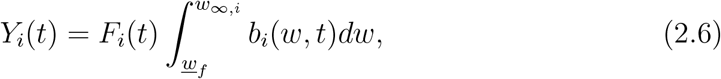

which becomes *Y*_*i*_(*t*) = *F*_*i*_(*t*)*B*_*i*_(*t*) when *B*_*i*_(*t*) is measured over the harvested size range.

Three kinds of fishing were considered:

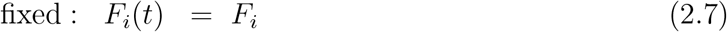

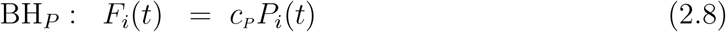

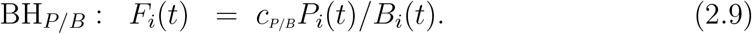

For the purpose of defining *F*_*i*_s, we set the body mass range 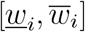 of *P*_*i*_(*t*) and *B*_*i*_(*t*) to match the harvested size range, i.e. *<u>w</u>i* = *<u>w</u>f* and 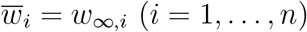. This is on the grounds that reliable information is most likely to be available over this range. Consistency of Eq. (2.8) requires *c*_*P*_ to have dimensions L2M^−1^. Rearranging Eq. (2.9), shows that the dimensionless constant *c*_*P/B*_ is equivalent to the exploitation ratio *E*_*i*_ = *Y*_*i*_*/P*_*i*_ for species *i*. In other words, *c*_*P/B*_ defines a fixed exploitation ratio that applies to all fish species. The exploitation ratio, also written as *E* = *F/M*, is widely used in fisheries science as a check on safe levels of fishing (Patterson, 1992), and is taken to indicate overfishing when above 0.5, or more conservatively when above 0.4 (Patterson, 1992; Pikitch et al., 2012).

### 2.4 Calibration

To study BH, we constructed model ecosystems from Eqs (2.1), (2.2), to capture some basic properties of real-life marine ecosystems. Key properties are as follows.

- It is well established that biomass *B* and production rate *P* of species are strongly positively correlated in marine ecosystems, and Eq. (2.4) reflects this. Fig. 2a, gives an example, using data from an Ecopath model of the West Scotland shelf ecosystem (Alexander et al., 2015), built as far as possible on observations of biomass and mortality rates, and assuming that biomass loss is balanced by production. This is in line with results from 110 Ecopath models of marine ecosystems summarised in Heath et al. (2017, Fig. BA2). Any study that intends to bring fishing of multiple species in line with *P* or *P/B* should respect this basic relationship.

**Figure 2:**
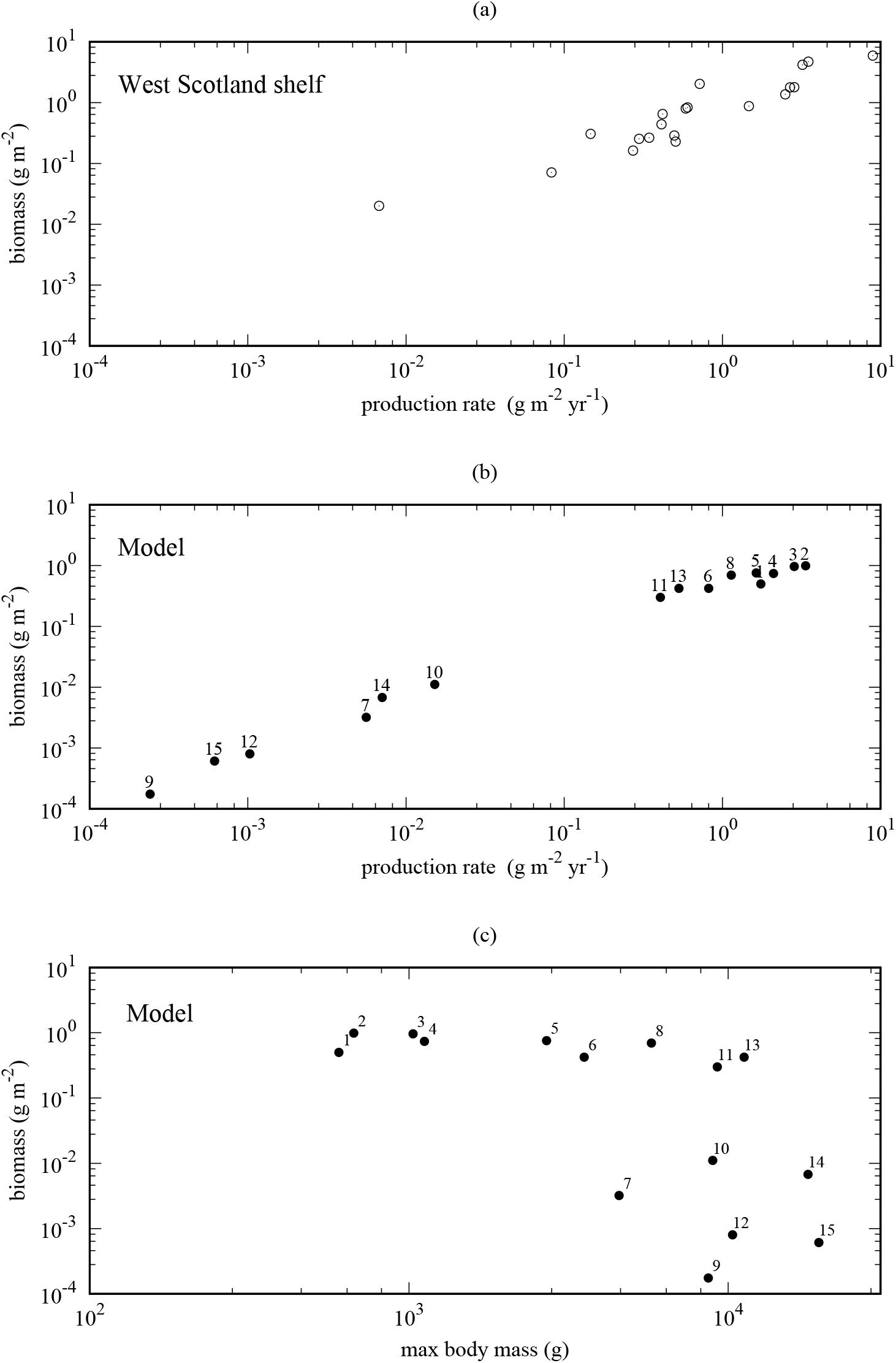
(a) The relationship between production rate and biomass from an Ecopath model of the West Scotland shelf ecosystem; open circles show values for individual species, with aggregated categories excluded. (b), (c) A model ecosystem with 15 unexploited fish species, supported by a plankton community. (b) Relationship between total biomass and total production rate of the modelled species close to equilibrium. Species numbered in rank order of *w*_∞,*i*_, as shown in (c).
- Ecological systems have a basic property that most species are rare (Baldridge et al., 2016). This applies to marine ecosystems as much as it does to others (Connolly et al., 2014). Both *B* and *P* fall to zero as abundance goes to zero, so there should be a tail of points for rare species going down to the bottom left corner of Fig. 2a. Ecopath models miss much of this tail, omitting most rare species, or aggregating them into broader categories: ‘rays’, ‘sharks’, ‘other small fish’ and ‘other pelagics’ in the West Scotland shelf model (Alexander et al., 2015). This paper is especially concerned with the effect of exploitation on rare species, because of the important contribution these species make to biodiversity. We therefore made sure that the model ecosystems would include rare species, despite their omission from empirical studies.
- Production rate of consumers is limited ultimately by the rate of primary production. To tie the model to real-world primary production, we calibrated it to Fig. 2 of San Martin et al. (2006), using the plankton carrying-capacity function *ã* (*x*), Eq. (2.2). This gave a total production rate of plankton of approximately 4000 g m^−2^ yr^−1^ (or t km^−2^ yr^−1^), roughly equivalent to 400 g carbon m^−2^ yr^−1^. This value is in accordance with the main cluster of observed values of primary production rate in a global analysis of Large Marine Ecosystems (LMEs) (Chassot et al., 2010, Fig. 1).
- Estimates of total yields from LMEs were also given by Chassot et al. (2010). The yields aggregated over species in our calibrated model ecosystems were in the range 0.2 → 1.0 g m^−2^ yr^−1^, consistent with the main cluster of values in Chassot et al. (2010, Fig. 1c: 0.2 → 1.0 t km^−2^ yr^−1^).
- The search-rate coefficient *A*_*i*_ of predators in Eq. (A.2) controls the rate at which fish accumulate body mass, and has usually been derived through a volume searched per unit time (Ware, 1978). It is dealt with differently here, because the measure of density is per unit area (not per unit volume). We calibrated *A*_*i*_ so that fish would grow from egg size to a realistic value of approximately 1 g in 1 yr through their feeding activities. We also checked the robustness to departures from this value by introducing an additional species-dependent random factor in *A*_*i*_ for the computations used in Figs 5 and 6 (Appendix B).

## 3 Results

To examine effects of fishing, we started by constructing unexploited model ecosystems. These provided baseline systems with the key empirical ecosystem properties listed above (Section 2.4 Calibration), to which fishing could be subsequently applied. The effects of different kinds of fishing on such ecosystems are described in detail in the first example below, and we follow this with a summary of three further examples, to demonstrate that the results have some generality and are not highly sensitive to a specific model ecosystem.

The ecosystems were constructed by sequentially adding fish species to plankton assemblages until 15 species were present (see Appendix B). (The upper limit on species kept computations manageable.) The species were picked with random maximum body masses *w*_∞,*i*_ and random larval mortality rates. The model ecosystems always captured the basic relationship between *B* and *P* observed in real-life ecosystems (Alexander et al., 2015) (Fig. 2a). We chose examplars that had tails of rare species that must exist in reality, even though they might often be missing from data collected for fisheries management. Our first example shows these features (Fig. 2b, c). The random *w*_∞,*i*_s of the fish species in this instance spanned a substantial part of the range of interest for exploitation, with a group of nine common species with *w*_∞,*i*_s from about 0.6 to 10 kg, and a group of six rarer species with *w*_∞,*i*_s from about 4 to 20 kg (Fig. 2c). We refer to the two groups as ‘common’ and ‘rare’ below. In the unexploited state, some species’ biomasses were still changing slowly after 50 years, so the state at this time is best thought of as a quasi-equilibrium.

Fig. 3 compares three contrasting regimes for exploiting the 15-species fish assemblage: fixed *F*, BH_*P*_, and BH_*P/B*_, as defined in Eqs (2.7)–(2.9). To show the differences between fishing regimes as simply as possible, we excluded variation in fishing intensity between species within fishing regimes here. This makes the basic similarities and differences between the fishing regimes clear, although some-what different from current fisheries practice built on maximum sustainable yields (MSYs). (See Discussion for issues about MSY in an ecosystem context). We introduce random variation between species in fishing intensities later (Fig. 6), as a partial check that the effects on rare species are robust. Harvesting was started initially with the ecosystem in its unexploited state (Fig. 2), and continued for 50 years. All species were harvested from a minimum body mass of 400 g upwards (i.e. 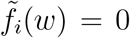 for *w <* 400 g). For comparability, the constants *c*_*P*_, *c*_*P/B*_ in Eqs (2.8), (2.9) were calibrated against a fixed fishing mortality rate *F*_*i*_ = *F* = 0.1 yr^−1^ Eq. (2.7), so that all three regimes would generate similar ecosystem biomass yields after 50 years, around 0.25 g m^−2^ yr^−1^. Specifically, BH_*P*_ took *F*_*i*_(*t*) = *c*_*P*_ *P*_*i*_(*t*) with *c*_*P*_ = 1 m2 g^−1^, and BH_*P/B*_ took *F*_*i*_(*t*) = *c*_*P/B*_*P*_*i*_(*t*)*/B*_*i*_(*t*) with *c*_*P/B*_ = 0.25. The ecosystem-level yield is consistent with values regarded as acceptable in large marine ecosystems around the world (Chassot et al., 2010; Link and Watson, 2019).

**Figure 3:**
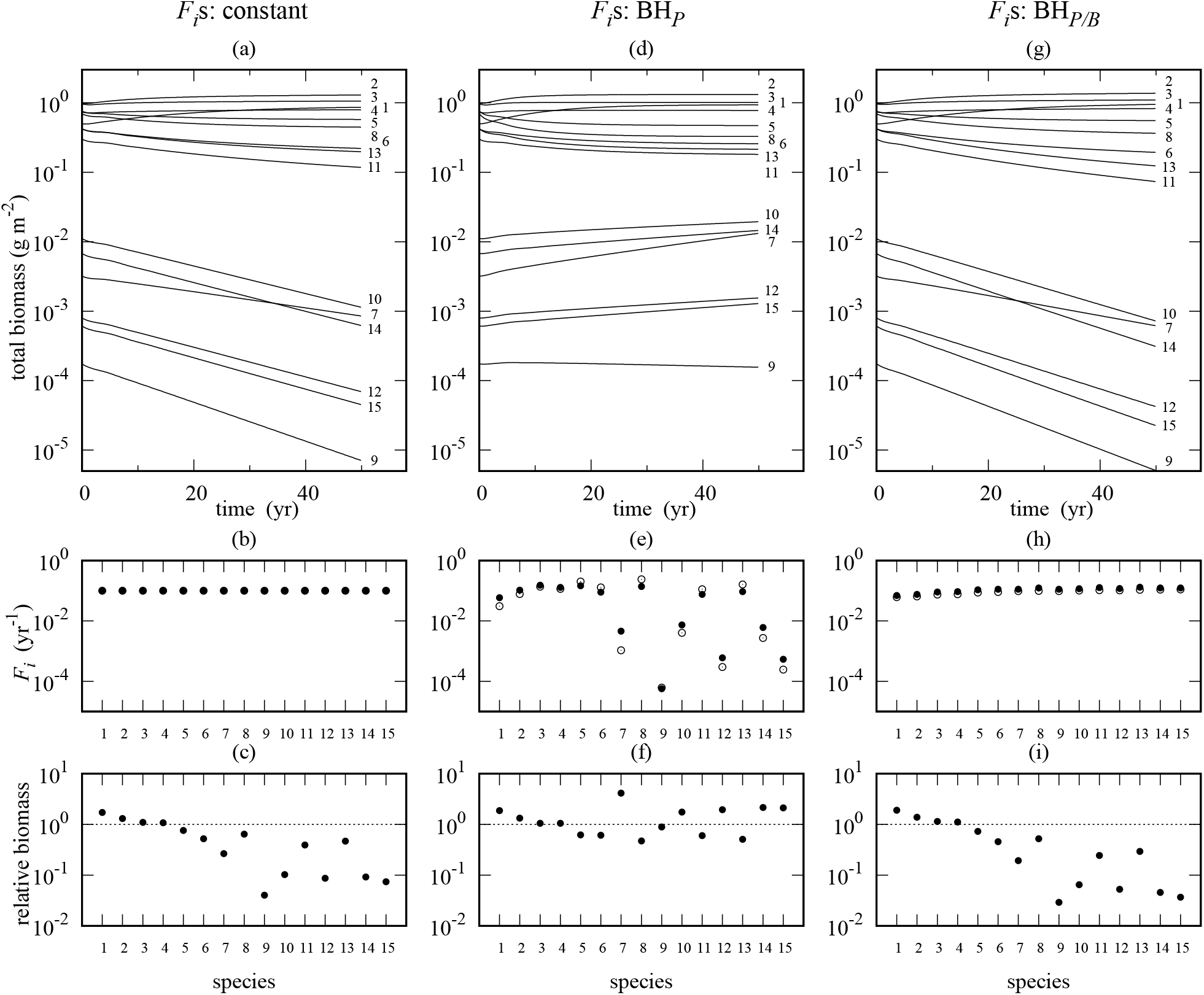
Three ways to harvest a multispecies fish assemblage. Fishing mortality rates (*F*_*i*_) were set: constant over time and species (column 1), adaptively using BH_*P*_ (column 2), adaptively using BH_*P/B*_ (column 3). Panels a, d and g show the changes in total biomass of species following the start of fishing in year 0. Panels b, e and h show the *F*_*i*_s. BH is adaptive, and allows *F*_*i*_ to change over time as the ecosystem changes: open circles in panels e and h show *F*_*i*_ in year 0, and filled circles show *F*_*i*_ in year 50. Panels c, f and i show the ratio of the total biomass in year 50 to the total biomass in the unexploited ecosystem in year 0 (Fig. 2b); points on the dotted lines (ratio = 1) would indicate no change in total biomass caused by fishing. Numbers 1 to 15 identify fish species in rank order of *w*_∞,*i*_, as given in Fig. 2c.

Fig. 3 shows that only BH_*P*_ supported the rare species (Fig. 3a, d, g). Moreover, BH_*P*_ differs from fixed *F*_*i*_ only in that the fishing mortality rates were weighted adaptively over time by the current somatic production rates (*P*_*i*_’s) in the exploited range of body sizes. The rare species were protected by their low production rates, which made their *F*_*i*_s much smaller than those of the common species (Fig. 3e). This weighting of fishing mortality rate prevented the decline in the rare species, and kept both the rare and common species quite close to their unexploited biomasses (Fig. 3f). (This is with the caveat that harvesting still truncated the size distribution of common species.) The *F*_*i*_s, being adaptive, allowed species to compensate to some extent for changes in the flow of mass over time caused by fishing, decreasing in species with biomass ratios less than one, and increasing in species with biomass ratios greater than one (Fig. 3e, f).

The third column of Fig. 3, BH_*P/B*_, weights the fishing mortality rates, *F*_*i*_, adaptively by the ratio *P*_*i*_(*t*)*/B*_*i*_(*t*). In this case, the constant *c*_*P/B*_ in Eq. (2.9) was set to give an exploitation ratio *E* = 0.25, well below the accepted maximum value for safe fishing (Patterson, 1992). This generated *F*_*i*_s near the constant value 0.1 yr^−1^ in Fig. 3b. So the outcome was similar to the fixed fishing strategy shown in column 1, with the rare species collapsing. Although the *F*_*i*_s could, in principle, adapt to changes in species abundance, in practice they were rather unresponsive to the decline of rare species (Fig. 3g), and remained close to those in column 1 (Fig. 3h).

The cause of the difference between the exploitation methods becomes clear from plotting the log-transformed yields and production rates (Fig. 4). In such plots, constant exploitation ratios *E* are isoclines with slope 1. Unsurprisingly, the main contribution to yield came from the high-biomass species in the top-right corner of the plots. From a fisheries perspective, the other species would likely just be a rare by-catch in a mixed-species fishery.

**Figure 4:**
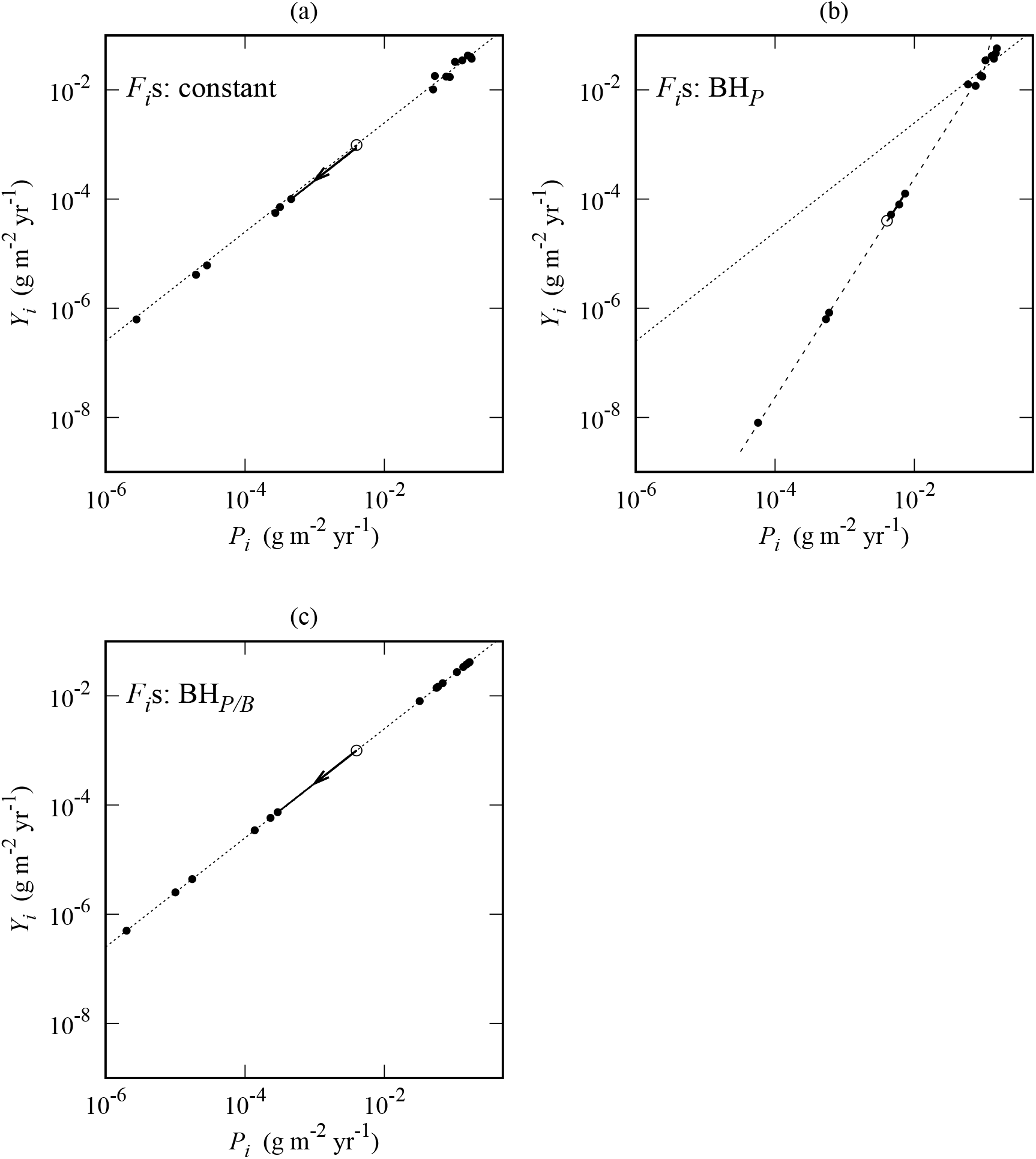
Yields and production rates from a multispecies fish assemblage exploited in three ways, as described in Fig. 3. Filled circles are values for 15 fish species after 50 years of harvesting, the species ordered as in Fig. 3. The species are approximately in balance near the dashed line in (b) with slope 1 + *α* = 2.004 (see text), but not near the dotted lines of constant exploitation ratio *E* = 0.25 in (a) and (c). As an example of the imbalance in (a) and (c), the trajectory of species 10 from year 0 (open circle) to year 50 is shown (the continuous line, with an arrow indicating the direction of change.) The range of body mass over which the production rate *P*_*i*_ was measured, Eq. (2.4), was set to match the harvested size range, i.e. from 400 g to *w*_∞,*i*_.

**Figure 5:**
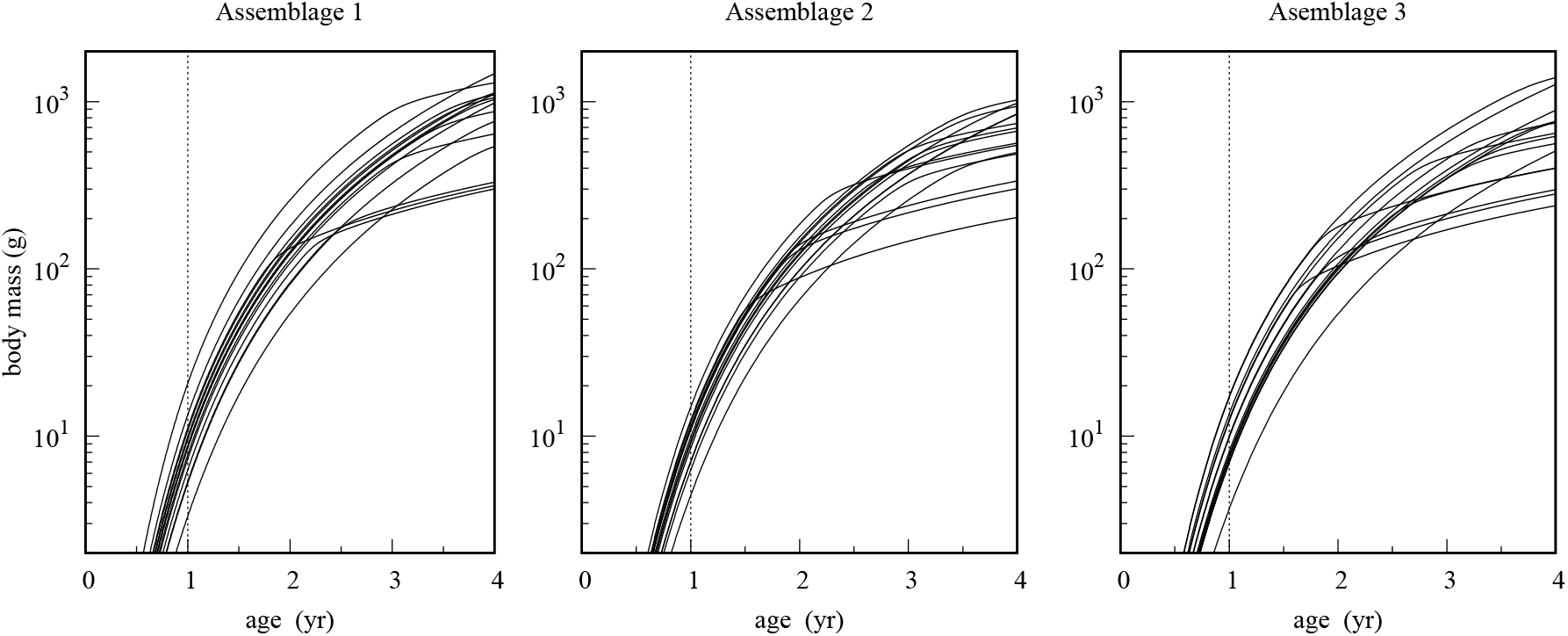
Trajectories of somatic growth of 15 species in three independent replicate assemblages, with random variation among species in food-search activity (Appendix B). These trajectories emerge solely from food eaten by the fish, and were obtained by solving Eq. (C.1) in Appendix C. The growth curves were measured when the assemblages were close to equilibrium in the absence of fishing.

**Figure 6:**
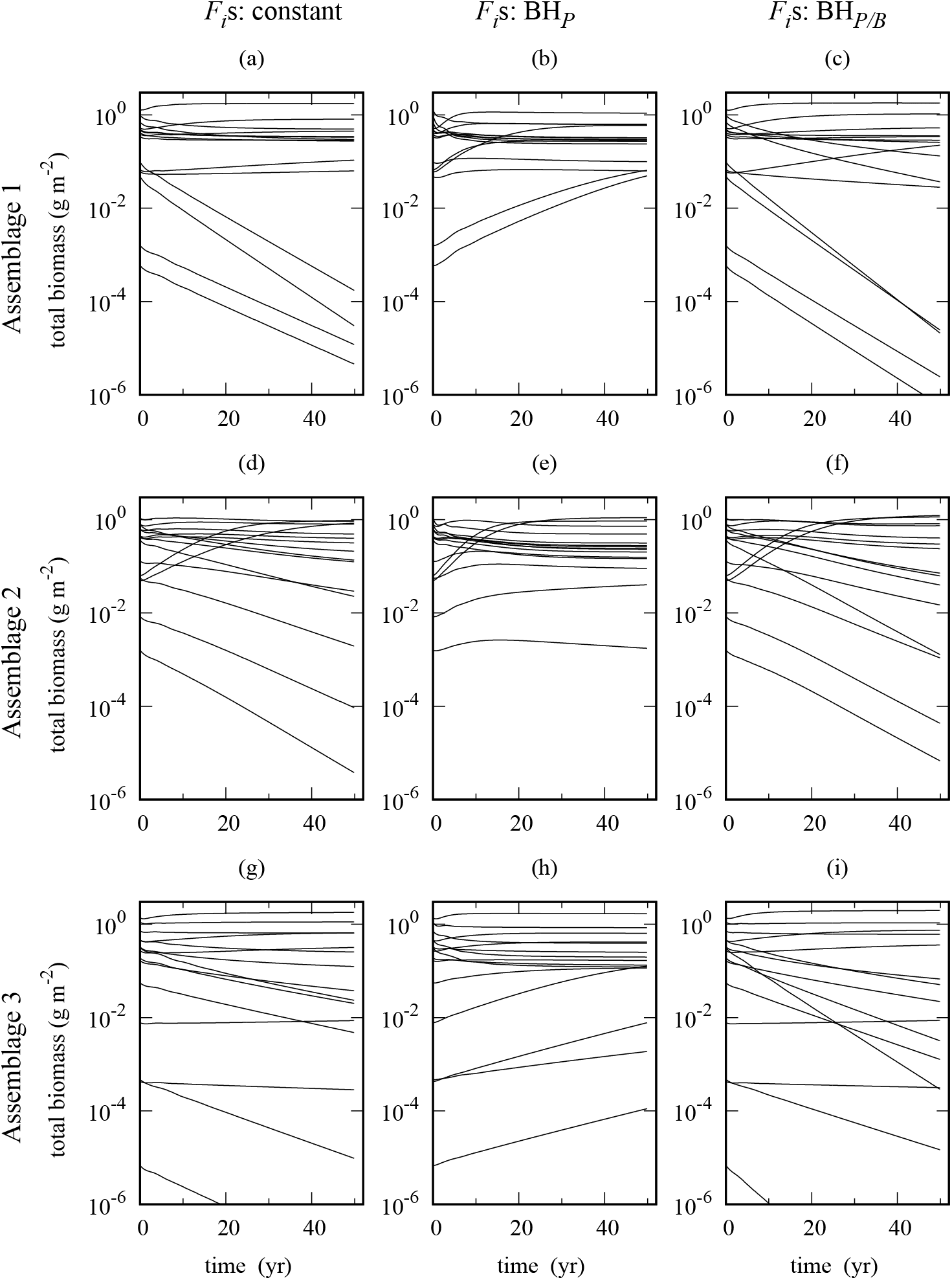
Tests of robustness of results in 3 independently-assembled ecosystems (rows 1,2,3), each containing 15 fish species. These assemblages contained greater life-history variation between species than the ecosystem used in Figs 2, 3, 4. Also, the baseline fishing mortality was doubled, and random variation between species in fishing intensities introduced. The randomisations are described in Appendix B. Fishing was applied for 50 years, starting in year 0, and all species were harvested from a minimum body mass of 400 g upwards. Fishing mortality rates (*F*_*i*_s) in column 1 (a,d,g) were constant over time, in column 2 (b,e,h) were set by BH_*P*_, and in column 3 (c,f,i) were set by BH_*P/B*_.

The rare species, although of little significance for yield, were very much affected by how harvesting was done. BH_*P/B*_ did exactly what it was designed to do, bringing the species onto an isocline of constant *E*, in this instance at *E* = 0.25 (Fig. 4c). But, importantly, the resulting assemblage was not in balance: the rare species simply followed a downward path along the line *E* = 0.25. To illustrate this, the trajectory of species 10 is plotted from year 0 to 50 in Fig. 4c. The other five rare species were collapsing in a similar way. (Constant *F*_*i*_ led to a similar collapse, as illustrated for species 10 in Fig. 4a.) The reason why BH_*P/B*_ allows this to happen is that biomass is a core component of production rate (see Eq. (2.4)), so *P*_*i*_ and *B*_*i*_ rise or fall together, while their ratio *P*_*i*_*/B*_*i*_ changes relatively little. So it is quite possible for species to be on a path to extinction, while the ratio *P*_*i*_*/B*_*i*_ remains close to constant. The ratio might even increase if low species biomass reduced competition for food. Like the fixed-fishing regime, BH_*P/B*_ is not a sensible strategy for maintaining biodiversity.

To maintain biodiversity calls for a different relation between yield and production (Fig. 4b). BH_*P*_ makes *Y*_*i*_ = *F*_*i*_*B*_*i*_ = *c*_*P*_ *P*_*i*_*B*_*i*_. Since *P*_*i*_ and *B*_*i*_ are closely related, this means yield should have a nonlinear relationship with production. Given a scaling relationship of the form 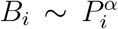 (e.g. Fig. 2a, b), the relation needed 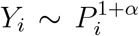. In the model ecosystem (Fig. 2b), *α* = 1.004 when based on the harvested body-size ranges, so the species were near a line of slope 2 in Fig. 4b. It was harvesting close to this line that kept the species in balance, protecting the rare species, and maintaining biodiversity, while giving a biomass yield similar to the other fishing regimes. The results in Figs 3, 4 therefore suggest that fishing for biodiversity calls for *F*_*i*_ ∝ *P*_*i*_, rather than *F*_*i*_ ∝ *P*_*i*_*/B*_*i*_. Put another way, species need increasing protection from fishing through a reduction in the exploitation ratio, as biomass (and hence production rate) goes down.

The results are potentially sensitive to assumptions about the life histories of the species. Somatic growth in particular plays a key role in the measure of production rate, Eq. (2.4), and had relatively little variation between species at early life stages in Figs 2, 3, 4. As a partial check on robustness of the results, we therefore assembled further independent ecosystems, with random variation among species in the rate at which they searched for food (the search-rate coefficient *A*_*i*_, see Appendix B).

This allowed fish which searched more actively to grow faster, the benefit of faster growth being offset by greater intrinsic mortality. Trajectories of somatic growth of species in three such assemblages are shown in Fig. 5. By age 1 year, body mass spanned a four- to seven-fold range; growth slowing down later on at maturation, as food was transferred increasingly to reproduction. We also treated fishing intensity as a random variable and, at the same time, doubled the baseline fishing mortality rate from *F*_*i*_ = 0.1 → 0.2 yr^−1^ (Appendix B). The effect of applying fishing to these assemblages is shown in Fig. 6, and leads to the same conclusion as before: fishing by BH_*P*_ can maintain biodiversity. Doubling the fishing intensity and introducing the additional variation in life histories and fishing mortality rates led to more change in the time series under BH_*P*_, and a less faithful match to the natural biomass of species in the ecosystems than in Fig. 3. Nonetheless, we have found no instance of exponential decline of species in fisheries under BH_*P*_, which was a recurring feature of constant fishing mortality rates, and of fishing mortality rates set by BH_*P/B*_. This is not surprising. BH_*P*_ makes fishing density-dependent, since the fishing mortality rate responds to changes in the production rates, which themselves depend heavily on biomass (Eq. (2.4)). Clearly, such density dependence was absent when *F*_*i*_s were held constant, and it was also largely absent when *F*_*i*_s were set by the ratio *P*_*i*_(*t*)*/B*_*i*_(*t*) in BH_*P/B*_.

## 4 Discussion

The results from this study suggest how fishing mortalities could be arranged across species, to help keep the structure of exploited fish assemblages intact, while allowing some exploitation. In particular, the results show that BH_*P*_ provides a scaling of fishing mortality to production rates that gives some protection to rare species, thereby helping to maintain biodiversity in fish assemblages. (The scaling of BH_*P/B*_ to a constant exploitation ratio *E* does not achieve this.) Such fishing mortality spans a much wider range than would normally be considered, with low-production species experiencing very low fishing mortality. In effect, the rarer the species are, the more protection from exploitation they need. In practical terms, abundant species need to be carefully selected for exploitation, to satisfy simultaneously the requirements of fisheries and conservation.

Setting fishing mortality in proportion to production rate (BH_*P*_) is not in general equivalent to setting a constant exploitation ratio *E*, which forms the basis of BH_*P/B*_. Equivalence would need *P* to be independent of *B*, i.e. a scaling parameter *α* = 0 in 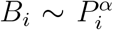. Real-world assemblages do not satisfy this condition. This is because they are characterised by a small number of common species and many less common ones (Connolly et al., 2014). Given the close formal link between biomass and production rate (Eqs (2.3), (2.4)), a strong, positive correlation between *P* and *B* should usually be the norm across species in marine assemblages, as is indeed widely reported in Ecopath models of marine ecosystems (see Heath et al., 2017, Fig. BA2). The value of *α* is an empirical matter, and it might well differ from one ecosystem to another, and even change over the course of time. But in any event, it would be surprising to find *α* = 0. Importantly, this means that empirical log*Y* –log*P* plots with slopes close to 1 (i.e. similar exploitation ratios across species), do not indicate good health of exploited ecosystems (Kolding et al., 2016). The evidence here is that fishing near a constant exploitation ratio does not bring an assemblage of fish species into balance. Even if this ratio is kept moderate, and even if the species start relatively abundant, such fishing can still put some species on a downward path.

The results suggest it could help to monitor the impact of exploitation at the ecosystem level through the relationship between fishing mortality and production species by species, irrespective of any debate about benefits of balanced harvesting (Froese et al., 2016; Pauly et al., 2016; Rehren and Gascuel, 2020). Monitoring can be done using estimates of fishing mortality rate, production rate and biomass, as they become available in real-time. Production rate is especially useful, as it automatically integrates over all the paths by which biomass flows into components of an ecosystem (such as a harvested body-mass range of a species). Production rate can be estimated from somatic growth rate and biomass (Heath et al., 2017), and can therefore substitute for detailed information on mass flow, when there is uncertainty about its path through an ecosystem, as there almost certainly will be. Such monitoring provides early feedback on ecosystem effects of fishing on commercial species, which is important given the complexity of marine ecosystems and our limited knowledge of how they work. (This information is less likely to be available for rare species.) At present, we know of only one published study of the relationship between fishing mortality and production rate, a report on the west of Scotland shelf ecosystem, using 1985 data in an Ecopath model (Alexander et al., 2015). It is notable that study found no relationship between fishing mortality and production rates of species in the ecosystem (Heath et al., 2017).

In fact, Eq (2.4) points to three separate reasons why somatic production rate might be low, of which this paper has considered just one, that of low biomass 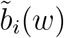. A second cause is that, as fish mature, incoming biomass is increasingly allocated to reproduction rather than to somatic growth, as given by the function 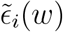. Fishing mortality from BH_*P*_, if disaggregated to distinguish juveniles from adults, can therefore be used to protect big old fish (BOFFFFs), and thereby facilitate the renewal of exploited populations (Hixon et al., 2014). A third cause is a shortage of food, giving a low mass-specific rate of food intake 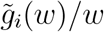. The effect of reducing fishing at a food-limited life stage is entirely different. It increases the biomass, further depleting the limited supply of food, generating a positive feedback in which somatic growth slows down even more. The outcome of this could be a population of stunted fish, like that of the Baltic cod (*Gadus morhua* L.) (Svedäng and Hornborg, 2017). This emphasizes the need for care in interpreting production rate. If this is falling when biomass of a species is large, scarcity of food is a likely cause, and potentially a signal of an underlying structural change in a marine ecosystem, which could be important to know about at this time of rapid climate change (Gaines et al., 2018).

Importantly, checks on production rates and biomasses of species, in their current ecosystem setting, are independent of information about an equilibrium, unexploited state of nature which would be needed to underpin a maximum sustainable yield (MSY). The scientific basis of MSY for single species has long been debated (Larkin, 1977; Finley and Oreskes, 2013), and becomes especially problematic in a multispecies setting, for several reasons. Knowledge of how these complex coupled systems operate is incomplete. Exploitation is likely to change the balance among species in ways that would be hard to unravel, and the shifting physical environment leaves its own footprint on the ecosystem. We know of no way to wind the clock back in these multispecies systems, to infer with precision what the biomasses of species would be in a natural state. From an ecosystem perspective, the unexploited biomasses of species, which form the basis of the MSY, are a matter of conjecture, and not a starting point for management. So, for instance, a recent suggestion that exploited species should be brought to around 60 % of their unexploited biomasses (Pauly and Froese, 2020) begs the question as to what these unexploited biomasses are.

Aligning fishing mortality to somatic production rate (BH_*P*_) can be thought of as a harvest control rule (Punt, 2010; Kvamsdal et al., 2016) for multiple species coupled together in ecosystems, as opposed to single species living in isolation. Its purpose is to set the relative fishing mortality rates across species so that rarer species get the protection they need, while allowing some exploitation of more common species. Counter-intuitive effects from fishing can readily emerge, once the coupling of species is allowed for. For instance, exploited piscivores have been seen to benefit from harvesting their forage fish (Soudijn et al., 2021), and we found cases where rare species gained more through release from predation under BH_*P*_ than they lost from fishing (results not shown). Making management responsive and adaptive helps to deal with unanticipated changes, and as would be expected, this worked better at lower, than at higher fishing intensities (compare Fig. 3d with Fig. 6b, e, h). The harvest control rule avoids the risk that fishing-induced deaths in rare species are seen as accidental collateral damage, about which little can be done. Instead, it places conservation on an equal footing with the fisheries that exploit an ecosystem.

We have not addressed the acceptable intensity at which to exploit marine ecosystems as a whole (the parameter *c*_*P*_ in Eq. (2.8)). This is a societal matter involving the public, together with stakeholders in fishing, conservation, markets and science (Heath et al., 2017). However, it is clear that conservation of marine ecosystems has to deal with the way in which fishing is distributed across species, as well as with the total ecosystem harvest (Link and Watson, 2019). A given total yield of biomass can be obtained with differing degrees of damage to biodiversity and ecosystem structure, depending on where that yield comes from (Figs 3, 4, 6). A moderate overall level of fishing is necessary, but how this fishing is distributed across ecosystem components is also a crucial question for meeting the needs of conservation of exploited marine ecosystems.

As in all modelling of complex systems, there are caveats to keep in mind. Size-spectrum models as used here, while removing some of the most serious limitations built into single-species fishery models, are still gross simplifications of marine ecosystems. The models assume ecosystems are non-seasonal, and the treatment of plankton dynamics is much simplified. The models assume the systems are well-mixed, and that organisms encounter other organisms in proportion to their spatially averaged abundance. Although the models do the bookkeeping of biomass as it flows from prey to predator, we have used simple assumptions about how this operates. Note also that the rare species of special importance for conservation, are also the species for which information on biomass, production rate and fishing mortality is most likely to be scarce. For simplicity, we picked a single illustrative pattern of fishing in which all species entered a mixed-species fishery at a single body mass, and there are clearly many other possibilities. In particular, we have not attempted to match fishing closely to the range of life histories in a natural fish assemblage. While the study can suggest how fishing mortality rates need to be set for conservation of biodiversity, there would be serious practical issues to resolve in making sure these rates are not exceeded in rare species. Moreover, the fish assemblage is only a subset of the species present, and we have not considered the many other species of special concern for conservation, such as seabirds, marine mammals, and benthic fauna, all of which can be heavily impacted by fishery decisions (Cury et al., 2011; Pikitch et al., 2012).

Obviously there is more to conservation of marine ecosystems than maintaining biodiversity. Aggregated measures of species biomass hide the truncation of size structures caused by fixed fishing mortality rates from recruitment onwards, and hide the strong directional selection on body size that can lead to fisheries-induced evolution (Law and Plank, 2018). Effective functioning of marine ecosystems calls for unconstrained flow of biomass from microscopic phytoplankton to large fish and mammals, and there is a need to understand how fishing can be organised to prevent fisheries-induced bottlenecks in the flow from building up (Svedäng and Hornborg, 2017). There is much to learn about how the footprint of fisheries on marine ecosystems can be reduced.

For all the limitations of modelling, aligning fishing mortality rate to production rate is intuitive, and the evidence is that it is not heavily model-dependent (see Plank, 2018). It points in a direction that could help reconcile fishing with some of the concerns of conservation. Given the increasing conflict between conservation and fisheries, the increasing restrictions being placed on fisheries, and the resulting damage to lives of those who depend on them, we suggest this route is worth considering.

## Acknowledgements

This work was initiated as an EPSRC IAA Strategic Partnerships Fund to the University of York UK for work at the Department of Mathematics and Statistics, University of Canterbury, NZ. RL was supported by the Natural Environment Research Council and Economic and Social Research Council funded Sustainable Management of UK Marine Resources (SMMR) programme (grant number NE/V01708X/1). MJP was supported by Te Pūnaha Matatini, a New Zealand Centre of Research Excellence in Complex Systems. We thank J. W. Pitchford for comments on the manuscript, and S. M. Garcia, J. Kolding, S. Zhou, with whom we have had extensive discussions about balanced harvesting.

## Data Availability Statement

Data sharing is not applicable to this article as no new data were created or analysed in this study.

## APPENDICES

### A Mathematical model

The ecosystems we studied span approximately 15 orders of magnitude in body mass from the smallest phytoplankton cells to the largest fish. We therefore transformed body mass *w* into a dimensionless logarithm, *x* = ln(*w/w*_0_) for the purpose of modelling. The constant *w*_0_ is an arbitrary body mass, taken as 1 g.

#### A.1 Fish dynamics

We write the state variable for fish species *i* as *u*_*i*_(*x, t*). Here *u*_*i*_(*x, t*)*dx* = *ϕ*_*i*_(*w, t*)*dw* is the density of individuals with log body mass in a small range [*x, x* + *dx*] at time *t*, per unit area of sea surface. The variable *u*_*i*_(*x, t*) is measured per unit sea surface area (dimensions: L^−2^), for consistency with the measure used for plankton production.

The dynamical system for *n* fish species is a system of *n* partial differential equations describing the flux in densities *u*_*i*_(*x, t*) (*i* = 1, …, *n*) generated by biological processes, given as:

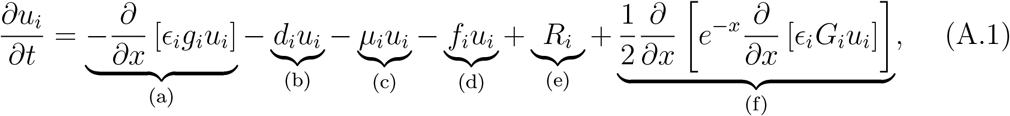

(Law et al., 2016). The function arguments *x* and *t* have been suppressed here and are given in full below. Eq. (A.1) equation is a second-order approximation to a jump-growth model, which itself is a systematic expansion of a master equation from a stochastic predator-prey process in which organisms grow and die through eating one another (Datta et al., 2010). The terms on the right-hand side of Eq. (A.1) describe: (a) somatic growth, (b) death from being eaten, (c) death from intrinsic causes, (d) death from fishing, (e) reproduction, (f) second-order diffusion. The lower bound of body size of species *i* in Eq. (A.1) is given by an egg size *x*_0,*i*_, and the upper bound by the size of the largest individuals *x*_∞,*i*_ defined in Eq. (A.8) below.

##### (a) Somatic growth

The function *g*_*i*_(*x, t*) is the mass-specific growth rate (dimensions: T^−1^) of species *i* at size *x* and time *t* from eating prey of all taxa, before partitioning incoming food into somatic growth and reproduction:

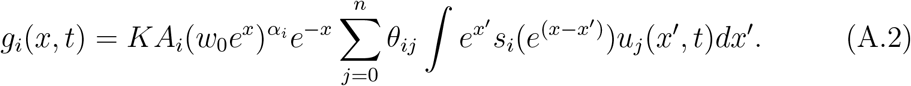

Here, the expression 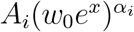 is an area over which a consumer of size *x* and species *i* feeds per unit time (dimensions: L2 T^−1^), where the term 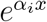 allows this area to scale with body size with allometric exponent *α*_*i*_. It would be more natural to use volume (Ware, 1978), but area is needed here because plankton production is measured per unit sea surface area. The constant *A*_*i*_ is a search-rate coefficient for species *i* (dimensions: L2 T^−1^ M^−*αi*^), and is calibrated to ensure realistic rates of individual growth. Dividing by the consumer size (*e*^−*x*^) makes the growth rate mass-specific. The integral is a convolution that adds up the gain in mass to the consumer of species *i* from eating items of species *j* (dimensions: L^−2^). This is a product of prey size 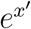, prey density *u*_*i*_(*x*^*′*^, *t*), and a dimensionless preference function for prey items of this size relative to the size of the consumer 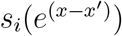, the product being integrated over prey sizes *x*^*′*^. The factor *θ*_*ij*_ allows different weightings of potential prey species. We adopt the notation that the first index refers to a predator, and the second index refers to a prey. The summation over *j* adds up the contribution made by all species to the growth of *i. K* is an efficiency with which prey biomass is converted into biomass of the consumer.

Expression (a) deals only with somatic growth, so *g*_*i*_(*x, t*) has to be multiplied by a dimensionless proportion of the food allocated to somatic growth *ϵ*_*i*_(*x*), Eq. (A.8) below. (The contribution of *g*_*i*_(*x, t*) to reproduction is dealt with separately in Eq. (A.9).) Multiplying by *u*_*i*_(*x, t*) converts the per capita rate into a population rate. The partial derivative ∂*/*∂*x*, known as an advective derivative, is needed because the change in density in the small size range [*x, x*+*dx*] depends on the rate at which density increases from growth into the range, and the rate at which density falls from growth out of the range.

##### (b) Death from being eaten

This is the flip side of growth, because it accounts for the feeding by other organisms on species *i* at size *x*. The function *d*_*i*_(*x, t*) is the per-capita death rate (dimensions: T^−1^) of species *i* at size *x* time *t*, due to all sources of predation:

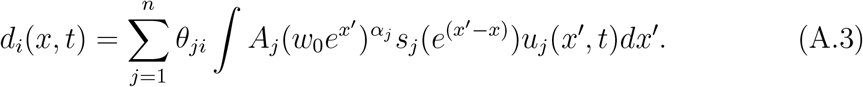

The integral is a convolution that integrates over body size *x*^*′*^ of predator species *j* to obtain its contribution to the per-capita predation rate on species *i* at size *x*, with symbols as defined in Eq. (A.2). The summation adds up the contributions of the different predator species, starting at *j* = 1 because plankton (represented by *j* = 0) are assumed not to eat fish, the parameter *θ*_*ji*_ being the weighting attached to predation by species *j* on species *i. θ*_*ji*_ has been interpreted as a measure of the spatial co-occurrence of species *i* and *j* (Blanchard et al., 2014; *Spence et al*., *2021*); *this has a symmetry θ*_*ji*_ = *θ*_*ij*_ for *i, j* ≥ 1 which we use here. The total rate at which density falls due to predation is then the per-capita rate *d*_*i*_(*x, t*) multiplied by the density *u*_*i*_(*x, t*).

###### Feeding kernel

The dimensionless function 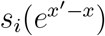 describes the preference of a consumer of size *x*^*′*^ and species *i* for prey of size *x*. It is placed inside the functions *g*_*i*_, *d*_*i*_, *G*_*i*_, all of which depend on predation. We use a simple box kernel, unselective on the log scale of body mass over a chosen range relative to the consumer:

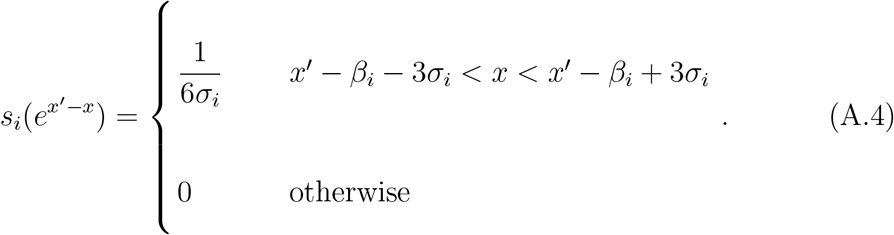

The factor of 1*/*6*σ*_*i*_ is a normalisation constant chosen such that the feeding kernel integrates to 1, i.e. 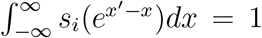. The function is given in this form so that *β*_*i*_ is the centre of the kernel, representing the mean log consumer:prey body mass ratio, and 6*s*_*i*_ is its width. This means that there is no feeding with a log consumer:prey body mass ratio outside the range [*β*_*i*_ − 3*s*_*i*_, *β*_*i*_ + 3*s*_*i*_].

##### (c) Death from intrinsic causes

We take the per-capita death rate from intrinsic causes *µ*_*i*_(*x, t*) (dimensions: T^−1^) in two parts, comprising a larval and a background component:

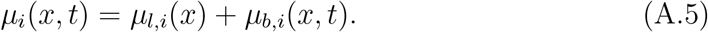

The total rate at which density falls due to these intrinsic processes is then the per-capita rate *µ*_*i*_(*x, t*) multiplied by density *u*_*i*_(*x, t*).

###### Larval mortality rate

We assume the extra risks of mortality at the larval stage are given by a per-capita larval mortality rate *µ*_*l,i*_(*x*) (dimensions: T^−1^) of species *i* at body size *x*:

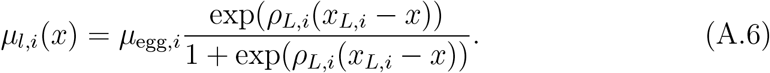

This is a reverse-sigmoid function that starts large on eggs, and falls to zero as the fish get big enough to leave the vulnerable stage. The function contains three parameters which can be species-dependent: (1) the magnitude of the extra mortality rate *µ*_egg,*i*_, as the larval stage begins, (2) the body size *x*_*L,i*_ at which fish are growing out of the vulnerable larval stage, and (3) the range of body size *ρ*_*L,i*_ over which this escape takes place. (Large *ρ*_*L,i*_ means this contribution to mortality declines sharply as fish leave the larval stage; small *ρ*_*L,i*_ means there is a more gradual transition from high to low larval mortality.)

###### Background mortality rate

The function *µ*_*b,i*_(*x, t*) (dimensions: T^− 1^) accounts for all intrinsic mortality other than the larval component. To set a level playing-field across species, we assume that this is proportional to the mass-specific needs for metabolism, relative to the mass-specific rate at which food becomes available at size *x*, based on a model in Law et al. (2016). These rates are set relative to their values at egg size, so 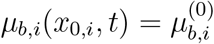 is a fixed baseline intrinsic mortality at birth for species *i*. The metabolic need should scale with body mass, and we write this as exp(−*ξ*(*x* − *x*_0,*i*_)), using the same exponent parameter *ξ* for all species. The mass-specific rate of food intake at size *x* relative to size *x*_0,*i*_ is *g*_*i*_(*x, t*)*/g*_*i*_(*x*_0,*i*_, *t*). Thus

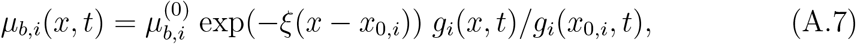

which is a function of time because it depends on the mass-specific growth rate *g*_*i*_(*x, t*).

##### (d) Fishing mortality rate

This depends on the rules chosen for fishing. We write this as a general per-capita death rate, *f*_*i*_(*x, t*) (dimensions: T^−1^) for species *i*, dependent on body size *x* and time *t*; the total rate at which density falls as a result of fishing is then its product with *u*_*i*_(*x, t*). The function *f*_*i*_(*x, t*) can have different forms, depending on the rules chosen for exploitation of the ecosystem.

##### (e) Reproduction

The dimensionless function 1 − *ϵ*_*i*_(*x*) describes the proportion of incoming food allocated to reproduction in species *i* at size *x*, and uses an expression suggested by Hartvig et al. (2011) and Law et al. (2012)

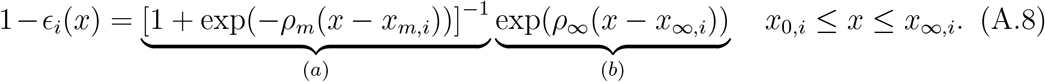

Part (a) deals with maturation, where *x*_*m,i*_ is the body size at which 50 % of the fish are mature in species *i*, and *ρ*_*m*_ defines the body-size range over which these fish are maturing. Part (b) describes allocation to reproduction once maturity is reached. The maximum body size *x*_∞,*i*_ of species *i* is typically very close to the size at which all incoming mass is allocated to reproduction and no further somatic growth is possible, the approach to *x*_∞,*i*_ being scaled by a parameter *ρ*_∞_.

The total rate *R*_*i*_(*t*) of egg production in species *i* at time *t* (dimensions: L^−2^ T^−1^) is then given by:

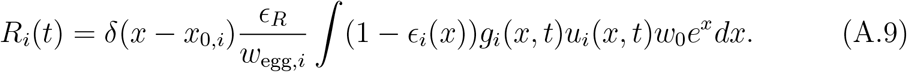

This takes the mass-specific rate at which reproductive mass is created at size *x*, multiplies by density and body mass to convert it to the total rate at which reproductive mass accumulates at body size *x*, and then integrates over all body sizes *x*. It then transforms the total mass rate into a rate of egg production by dividing by the egg mass *w*_egg,*i*_, allowing for some inefficiency (*ϵ*_*R*_ *<* 1) in converting the reproductive mass into eggs. The term *δ* (*x*−*x*_0,*i*_) is a Dirac delta function needed so that eggs enter the spectrum of species *i* in Eq. (A.1) only at the egg size *x*_0,*i*_.

##### (f) Diffusion

We included a second-order diffusion term (dimensions: L^−2^ T^−1^), to allow growth trajectories of fish to spread as the fish got older. Otherwise, all fish born at the same time would have identical trajectories of growth (Datta et al., 2010). The diffusion term contains a function

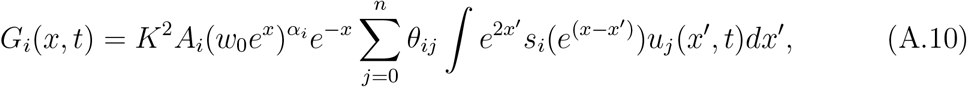

which is similar to the mass-specific growth rate in Eq. (A.2), where the terms it contains are defined.

#### A.2 Plankton dynamics

The plankton assemblage, which in reality could be of great complexity, was simplified to a single density function *u*_0_(*x, t*). Here *u*_0_(*x, t*)*dx* = *ϕ*_0_(*w, t*)*dw* is the density of individuals with log body mass in a small range [*x, x* + *dx*] at time *t*, per unit area of sea surface (dimensions: L^−2^).

The dynamical system for this aggregated plankton spectrum was written as a single partial differential equation operating at every body size *x*:

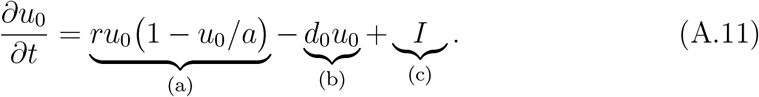

The terms on the right-hand side of this equation describe: (a) a core logistic function that drives the plankton dynamics, (b) a death rate from planktivory in the fish assemblage, and (c) an immigration term. There is no process of body growth here, so there is no direct coupling from one body size to another (in contrast to Eq. (A.1)). This assumption was made on the grounds that the plankton would be unicellular over most of the size range considered. The lower bound of body size in the plankton in Eq. (A.11) is given by the smallest cell size *x*_0,0_, and the upper bound by the size of the largest individuals *x*_∞,0_. Typically, size-spectrum models have used a linear semi-chemostat assumption in plankton to combine local dynamics and immigration (Hartvig et al., 2011). Eq. (A.11) makes these processes explicit.

To generate a realistic size spectrum for the aggregated plankton assemblage, the intrinsic rate of increase *r*(*x*) (dimensions: T^−1^), the carrying capacity *a*(*x*) (L^−2^), and the immigration rate *I*(*x*) (L^−2^ T^−1^) were set to scale with body mass 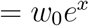 as:

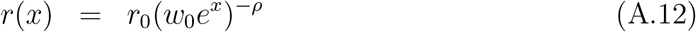

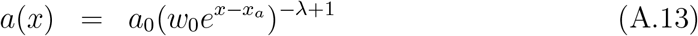

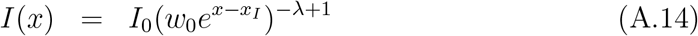

The function *r*(*x*) gives the cell division rate a scaling with cell size with an exponent *ρ* − 1 (Marañón et al., 2013), with the parameter *r*_0_ setting the overall rate. The function *a*(*x*) describes the carrying capacity the plankton would have in the absence of immigration and planktivory. This ensures that the size spectrum would settle to an equilibrium scaling with cell size with exponent −*λ* + 1 in the absence of other processes. The parameters *a*_0_ and *x*_*a*_ allow a calibration of the plankton spectrum to be made in line with natural plankton size spectra. The function *I*(*x*) was assumed to scale with body size in the same way as the carrying capacity. The function *d*_0_(*x, t*) is the per-capita death rate at size *x* and time *t* caused by planktivory in the fish assemblage, as given by Eq. (A.3) with *i* = 0.

### B Assembly and parameter randomisations

This work needed baseline ecosystems containing multiple size-structured species populations able to persist over the time scales needed for fishing. This is a subject of interest in its own right, because species coexistence is a core problem of community ecology. Here we outline how we constructed the ecosystems, as our approach was different from those used to construct multispecies size spectra in the past.

Multispecies size-spectrum models have typically added extra stock-recruitment relationships to species to achieve coexistence (e.g. Blanchard et al., 2014; Scott et al., 2014). However, the density-dependent feedbacks needed to generate stock-recruitment relationships are already built into size-spectrum models, and these relationships emerge naturally in a deterministic setting (Canales et al., 2020). Here we simply used the internal feedbacks in the models, avoiding the need to add arbitrary stock-recruitment relationships, and did three things to facilitate coexistence.

1. We included a key feedback (Ricker and Foerster, 1948; MacCall, 1980; Canales et al., 2020), which operates at the larval stage in the following way. When larval density is low relative to plankton food, the abundance of food allows fast body growth through the vulnerable larval stage, making the risk of death relatively small. Conversely, when larval density is relatively high, growth is slower and the accumulated risk of death in the larval stage becomes greater. This has a strong stabilizing effect when plankton are coupled to single-species size-spectra of fish (Canales et al., 2020, Fig. 2). It only requires mortality rate to be falling with increasing body size early in life, which is consistent with the results of the majority of studies (Sogard, 1997, Table 1). Food-dependent body growth, standardly built into size-spectrum models, does the rest.

**Table 1:** Plankton parameters.
2. The *θ*-matrix was set to reflect the qualitative structure of *θ*-matrices used in size-spectrum models of the North Sea (Blanchard et al., 2014) and Celtic Sea (Spence et al., 2021). These matrices were constructed from the spatial overlap of fish species, and have a property that diagonal elements are larger relative to off-diagonal ones, because species tend to show some spatial separation. The effect is to make cannibalism a stronger force relative to predation between species. Diagonal dominance is known to promote coexistence in simpler Lotka-Volterra food webs (Hofbauer and Sigmund, 1988, p. 193).
3. The ecosystems were constructed in steps, starting from plankton, adding one fish species at a time, leaving time for the system to relax before adding the next species. This reflects the ecological reality that multispecies systems do not spring into existence in their final form, but come together gradually as new species arrive, sometimes establishing themselves, and sometimes causing loss of species already present (Sigmund, 1995). Sequential assembly allows some sorting to take place, and is known to lead efficiently to the special sets of species that can persist in unstructured Lotka-Volterra systems (Lewis and Law, 2007).

| parameter | value | interpretation |
| --- | --- | --- |
| $w_0 e^{x_{0,0}}$ | $10^{-10}$ g | minimum body mass |
| $w_0 e^{x_{\infty,0}}$ | 1 g | maximum body mass |
| $w_0 e^{-x_a}$ | 0.001 g | calibrates $a(x)$ at 1 mg |
| $w_0 e^{-x_I}$ | 0.001 g | calibrates $I(x)$ at 1 mg |
| $r_0$ | $10 \text{ g}^{1-\rho} \text{ yr}^{-1}$ | sets intrinsic rate to $10 \text{ yr}^{-1}$ at 1 g |
| $a_0$ | $2000 \text{ m}^{-2} \text{ g}^{\lambda-1}$ | sets ‘equilibrium’ density to $2000 \text{ m}^{-2}$ at 1 mg |
| $I_0$ | $2000 \text{ m}^{-2} \text{ g}^{\lambda-1} \text{ yr}^{-1}$ | sets immigration rate to $2000 \text{ m}^{-2} \text{ yr}^{-1}$ at 1 mg |
| $\rho$ | 0.15 | scales intrinsic rate with respect to $x$ |
| $\lambda$ | 2 | scales slope of an ‘isolated’ size spectrum |

The term ‘equilibrium’ refers to an isolated plankton assemblage at cell size 1 mg. The term ‘isolated’ refers to a plankton spectrum lacking immigration and predation.

The algorithm for assembly started from an ecosystem containing plankton only, with densities close to the plankton carrying capacity function *a*(*x*) Eq. (A.13), and ran as follows.

1. Introduce one fish species, with log maximum body size uniformly distributed over [log(100*/w*_0_) ≤ *x*_∞,*i*_ ≤ log(40000*/w*_0_)] (i.e. maximum body mass is between 100 g and 40 kg), and larval mortality uniformly distributed over [28 ≤ *µ*_egg,*i*_ ≤ 32] yr^−1^, other parameters of the life history being held constant. The range for larval mortality was kept small, as species with high larval mortality tended to be excluded as the number of fish species increased. The size spectrum was started at a low density on a power law, the egg density being 0.002 m^−2^.
2. Allow the augmented ecosystem to relax towards an asymptotic state, by integrating Eqs (A.1), (A.11) over a 50 yr time period. Since some species could still be changing slowly after 50 yrs, the resulting ecosystems were no more than quasi-equilibrial.
3. After integration, remove all fish species with a total biomass *<* 2 *×* 10^−6^ g m^−2^. This extinction threshold was sufficient to allow an assemblage of fish species to build up, in which species biomasses spanned approximately four orders of magnitude.
4. If the number of fish species was *<* 15, and if the number of iterations was *<* 40, return to step 1. Otherwise end the assembly. The limit of 15 fish species was a compromise between the richness of real-world ecosystems, and a manageable computation time. The limit on the number of iterations was also to keep the computation time manageable. The limit on iterations also had the effect of excluding some instances of increasing resistance to invasion during assembly, accompanied by a sequence of species replacements, leading towards lower larval mortality.

From the output of the assembly algorithm we selected an ecosystem with 15 fish species for harvesting, satisfying the following criteria before fishing started. (1) The fish species should be able to coexist over a time period of 50 years. (2) The set of species should tend to a state close to equilibrium. (3) The species should have a wide range of maximum body mass within the limits 100 g to 40000 g. (4) The species should span approximately 4 orders of magnitude in biomass, because fishing for biodiversity would need to operate in ecosystems that contained both common and rare species. The algorithm was used to obtain the ecosystem analysed in Figs 2, 3 and 4, which reached the required number of fish species in 25 iterations.

For Figs 5 and 6, we introduced more life-history variation between species by making the search-rate coefficient of species *i* (parameter *A*_*i*_ in Appendix Eq. (A.2)) a random variable, *A*_*i*_*z*_*i*_, where *z*_*i*_ is a random draw from a normal distribution, *N* (*µ* = 1, *σ* = 0.1). Each search rate was offset by applying the same factor *z*_*i*_ to intrinsic mortality Eq. (A.5) *µ*_*i*_(*x, t*)*z*_*i*_ in species *i* (both the larval and background components). This created a trade-off in which species that searched more actively for food (and which therefore had faster somatic growth) also experienced a greater risk of mortality. The number of invasion attempts needed to create the 15-species assemblags was respectively: 24 (assemblage 1), 19 (assemblage 2), 45 (assemblage 3). (We allowed several extra iterations in assemblage 3, as it was close to 15 species by iteration 40.)

For Fig. 6, we also introduced random variation between species in the intensity of fishing and doubled the baseline fishing mortality from 0.1 to 0.2 yr^−1^. The *F*_*i*_s contained a species-dependent random factor, 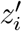, uniformly distributed over a range 0.5 to 1.5, and held constant over time. For comparability the same random factor was applied to all three methods of fishing within an assemblage. Thus, under constant *F*_*i*_, Eq. (2.7) was 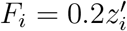 under BH_*P*_, Eq. (2.8) was 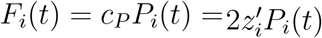 under BH_*P/B*_, Eq. (2.9) was 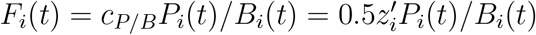.

### C Numerics

See Appendix B for the protocol used to generate random life histories. Other parameters in Eqs (A.1) and (A.11) were fixed and are described below.

Parameter values for plankton dynamics, Eq. (A.11), are given in Table 1. The lower bound of *x*, was set near the boundary of micro- and nanoplankton at 10^−1^0 g. A scaling relationship for cell division rate with cell size, as in Eq. (A.12), with an exponent near −*ρ* = −0.15 has been described starting near this cell size (Marañón et al., 2013). The upper bound of *x* was set at 1 g, to compensate for the absence of multicellular zooplankton in the model. Although these assumptions are artificial, they gave a total rate of primary production in keeping with observed values, after calibrating the carrying-capacity function of the plankton size spectrum, as described below.

The plankton carrying-capacity function *a*(*x*) was located at 2000 m^−2^ at a cell size 1 mg (using *a*_0_ = 2000, *x*_*a*_ = log(0.001)) from Fig. 2 of San Martin et al. (2006). The rate of primary production at size *x* was taken as 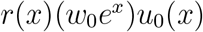, and the total rate as the integral of this over the range [*x*_0,0_, *x*_∞,0_]. On this basis, the total production rate of plankton was the region 4000 g m^−2^ yr^−1^ (or t km^−2^ yr^−1^), roughly equivalent to 400 g carbon m^−2^ yr^−1^. This is near the main cluster of observed values of primary production rate given in Chassot et al. (2010, Fig. 1). In Figs 3, 4, this production rate supported a total yield from the fish assemblage of about 0.25 g m^−2^ yr^−1^. Values given by Chassot et al. (2010, Fig. 1c), tended to be larger than this, but fishing was deliberately kept to a moderate level here. Our Fogarty ratio was about 0.06 ‰ in Figs 3, 4, well below 1 ‰, suggested by Link and Watson (2019) as the threshold for safety. The greater fishing intensities in Fig. 6 took the ratio up to 0.2 ‰, still well below the threshold. This suggests that levels of exploitation deemed safe for an ecosystem, could at the same time be unsafe for the species assemblages that live in them.

Parameter values for dynamics of fish species, Eq. (A.1), are given in Table 2. Life histories of fish species were constructed from a template common to all species (Fig. 1), with randomisation of certain parameters to generate differences between species (Appendix B). The template had a lower bound corresponding to egg mass, which was fixed for all species at 1 mg, and a random upper bound, as defined in Appendix B.

**Table 2:** Fish parameters.

| parameter | value | interpretation |
| --- | --- | --- |
| $w_0e^{x_{0,i}}$ | 0.001 g | egg mass |
| $w_0e^{x_{\infty,i}}$ | random g | maximum body mass |
| $\beta_i$ | 6.908 | centre of feeding kernel |
| $\sigma_i$ | 1.535 | measure of the width of feeding kernel |
| $A_i$ | $37.5 \text{ m}^2 \text{ yr}^{-1} \text{ g}^{-\alpha}$ | search-rate coefficient |
| $\alpha$ | 0.85 | search-rate scaling exponent |
| $K$ | 0.1 | ecological conversion efficiency |
| $\mu_{\text{egg},i}$ | random $\text{yr}^{-1}$ | larval death rate at egg size |
| $w_0e^{x_{L,i}}$ | 0.1 g | size around which larval death $\rightarrow 0$ |
| $\rho_{L,i}$ | 5 | exponent scaling size range as larval death $\rightarrow 0$ |
| $\mu_{b,i}^{(0)}$ | $0.1 \text{ yr}^{-1}$ | background death rate at birth |
| $\xi$ | 0.15 | exponent scaling falling death rate with size |
| $w_0e^{x_{m,i}}$ | random g | body mass at 50 % maturity: $w_0e^{x_{\infty,i}}/10$ |
| $\rho_m$ | 15 | exponent scaling size range of maturation |
| $\rho_{\infty}$ | 0.2 | exponent scaling approach to $x_{i,\infty}$ |
| $\epsilon_R$ | 0.2 | reproductive efficiency |

$A_i$ was randomised in Figs 5 and 6 (see Appendix B).

We used a box feeding kernel, in which the fish took prey items unselectively over a range 1/100000 to 1/10 of their own body mass. For instance, a newborn larval fish at 10^−3^ g would take prey items over a range [10^−8^, 10^−4^] g. The search-rate coefficient *A*_*i*_ has usually been derived through a volume searched per unit time, but is dealt with differently here, because the measure of density is per unit area (not per unit volume). We calibrated it so that fish would grow from egg size to 1 g in approximately 1 yr, with an additional random factor in the computations for Fig. 6 (see Appendix B).

Larval mortality rate *µ*_egg,*i*_ was set to a large value at birth, with fish leaving the larval stage *x*_*L,i*_ near to 0.1 g. After escaping from the larval stage, only a background mortality rate dependent on food intake remained. This was set by *µ*_*b,i*_ (0) to be small at birth, because of the explicit presence of larval mortality. In the computations for Fig. 6, an additional random factor was applied to these intrinisc components of mortality (Appendix B).

Following a commonly observed feature of fish life histories (Beverton, 1992; Froese and Binohlan, 2000), we assumed maximum body mass and mass at maturation were related. Thus maturation was also a random variable, taken as 1/10 of the random maximum body mass (see Appendix B).

A baseline of weights for predator-prey interactions in the ecosystem was set by an (*n*+1)*×*(*n*+1) matrix *θ*, with rows and columns indexed 0, 1, …, *n*, corresponding to plankton (*i* = 0) and *n* fish species (*i* = 1, …, *n*). Column 0 had *θ*_*i*,0_ = 1, for *i* = 1, …, *n*, meaning that all fish species could eat plankton. Row 0 had *θ*_0,*j*_ = 0, for *j* = 0, …, *n*, meaning that plankton could not feed on fish. The submatrix with rows and columns indexed 1, …, *n* described the fish assemblage, and had diagonal elements *θ*_*ii*_ = 0.5 for cannibalism, and off-diagonal elements *θ*_*ij*_ = 0.2 for *i*≠ *j*. This made self-limitation stronger than predation between species, giving some diagonal dominance to promote coexistence.

Eqs (A.1), (A.11) have to be discretised for numerical integration. We took a body-size step *δx* = 0.1, and a time step *δt* = 0.002 and used the Euler method. The speed of computation was increased by applying fast Fourier transforms to the convolution integrals in Eqs (A.2), (A.3), (A.10).

Growth trajectories were obtained by numerical solution of the differential equation:

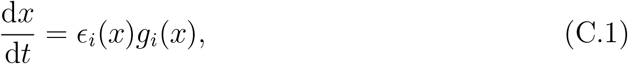

starting from an initial condition set by the egg size *x*(0) = *x*_0,*i*_.

## REFERENCES

Alexander, K. A., Heymans, J. J., Magill, S., Tomczak, M. T., Holmes, S. J., and Wilding, T. A. (2015). Investigating the recent decline in gadoid stocks in the west of Scotland shelf ecosystem using a foodweb model. ICES Journal of Marine Science, 72:436–449. doi:10.1093/icesjms/fsu149.

Allen, K. R. (1971). Relation between production and biomass. Journal of the Fisheries Research Board of Canada, 28:1573–1581.

Andersen, K. H. and Beyer, J. E. (2006). Asymptotic size determines species abundance in the marine size spectrum. American Naturalist, 168:54–61, doi:10.1086/504849.

Baldridge, E., Harris, D. J., Xiao, X., and White, E. P. (2016). An extensive comparison of species-abundance distribution models. PeerJ, 4:e2823:doi:10.7717/peerj.2823.

Beverton, R. J. H. (1992). Patterns of reproductive strategy parameters in some marine teleost fishes. Journal of Fish Biology, 41 (Supplement B):137–160.

Blanchard, J. L., Andersen, K. H., Scott, F., Hintzen, N. T., Piet, G., and Jennings, S. (2014). Evaluating targets and trade-offs among fisheries and conservation objectives using a multispecies size spectrum model. Journal of Applied Ecology, 51:612–622. doi:10.1111/1365–2664.12238.

Caddy, J. F. and Sharp, G. D. (1986). An ecological framework for marine fishery investigations. Fisheries Technical Paper 283, 152pp, FAO, Rome,Italy.

Canales, T. M., Delius, G. W., and Law, R. (2020). Regulation of fish stocks without stock–recruitment relationships: The case of small pelagic fish. Fish and Fisheries, 21:857–871. doi:10.1111/faf.12465.

Chassot, E., Bonhommeau, S., Dulvy, N. K., Mélin, F., Watson, R., Gascuel, D., and Le Pape, O. (2010). Global marine primary production constrains fisheries catches. Ecology Letters, 13:495–505. doi:10.1111/j.1461–0248.2010.01443.x.

Connolly, S. R., MacNeil, M. A., Caley, M. J., Knowlton, N., Cripps, E., Hisano, M., Thibaut, L. M., Bhattacharya, B. D., Benedetti-Cecchi, L., Brainard, R. E., Brandt, A., Bulleri, F., Ellingsen, K. E., Kaiser, S., Kröncke, I., Linse, K., Maggi, E., O’Hara, T. D., Plaisance, L., Poore, G. C. B., Sarkar, S. K., Satpathy, K. K., Schückel, U., Williams, A., and Wilson, R. S. (2014). Commonness and rarity in the marine biosphere. Proceedings of the National Academy of Sciences, 111:8524–8529. doi:10.1073/pnas.1406664111.

Cury, P. M., Boyd, I. L., Bonhommeau, S., Anker-Nilssen, T., Crawford, R. J. M., Furness, R. W., Mills, J. A., Murphy, E. J., Österblom, H., Paleczny, M., Piatt, J. F., Roux, J.-P., Shannon, L., and Sydeman, W. J. (2011). Global seabird response to forage fish depletion—one-third for the birds. Science, 334:1703–1706. doi:10.1126/science.1212928.

Datta, S., Delius, G. W., and Law, R. (2010). A jump-growth model for predator-prey dynamics: derivation and application to marine ecosystems. Bulletin of Mathematical Biology, 72:1361–1382. doi 10.1007/s11538–009–9496–5.

de Kerckhove, D. T. (2015). Promising indicators of fisheries productivity for the fisheries protection program assessment framework. Research Document 2014/108, DFO Canadian Science Advisory Secretariat Research Documnet, 200 Kent Street, Ottawa.

Dickie, L. M. (1972). Food chains and fish production. In Symposium on Environmental Conditions in the Northwest Atlantic, 1960-69, page 201–219. International Commission for the Northwest Atlantic Fisheries, special publication No. 8, Dartmouth, N.S., Canada.

Finley, C. and Oreskes, N. (2013). Maximum sustained yield: a policy disguised as science. ICES Journal of Marine Science, 70:245–250. doi:10.1093/icesjms/fss192.

Fowler, C. W. (1999). Management of multi-species fisheries: from overfishing to sustainability. ICES Journal of Marine Science, 56:927–932.

Froese, R. and Binohlan, B. (2000). Empirical relationships to estimate asymptotic length, length at first maturity and length at maximum yield per recruit in fishes, with a simple method to evaluate length frequency data. Journal of Fish Biology, 56:758–773. doi:10.1006/jfbi.1999.1194.

Froese, R., Walters, C., Pauly, D., Winker, H., Weyl, O. L. F., Demirel, N., Tsikliras, A. C., and Holt, S. J. (2016). A critique of the balanced harvesting approach to fishing. ICES Journal of Marine Science, 73:1640–1650. doi:10.1093/icesjms/fsv122.

Gaines, S. D., Costello, C., Owashi, B., Mangin, T., Bone, J., García Molinos, J., Burden, M., Dennis, H., Halpern, B. S., Kappel, C. V., Kleisner, K. M., and Ovando, D. (2018). Improved fisheries management could offset many negative effects of climate change. Science Advances, 4:doi:10.1126/sciadv.aao1378.

Garcia, S., Zerbi, A., Aliaume, C., Do Chi, T., and Lasserre, G. (2003). The ecosystem approach to fisheries. Issues, terminology, principles, institutional foundations, implementation and outlook. Technical Report FAO Fisheries Technical Paper. No. 443. 71 p, FAO, Rome.

Garcia, S. M., Kolding, J., Rice, J., Rochet, M.-J., Zhou, S., Arimoto, T., Beyer, J. E., Borges, L., Bundy, A., Dunn, D., Fulton, E. A., Hall, M., Heino, M., Law, R., Makino, M., Rijnsdorp, A. D., Simard, F., and Smith, A. D. M. (2012). Reconsidering the consequences of selective fisheries. Science, 335:1045–1047.

Garcia, S. M., Rice, J., and Charles, A. (2014). Governance of marine fisheries and biodiversity conservation: A history. In Garcia, S. M., Rice, J., and Charles, A., editors, Governance of Marine Fisheries and Biodiversity Conservation: Interaction and Coevolution, pages 3–17. John Wiley and Sons Ltd, Chichester UK.

Hartvig, M., Andersen, K. H., and Beyer, J. E. (2011). Food web framework for size-structured populations. Journal of Theoretical Biology, 272:113–122. doi:10.1016/j.jtbi.2010.12.006.

Heath, M., Law, R., and Searle, K. (2017). Scoping the background information for an ecosystem approach to fisheries in scottish waters: Review of predatorprey interactions with fisheries, and balanced harvesting. Technical Report FIS013, Fisheries Innovation Scotland. ISBN: 978-1-911123-10-1, available at: http://www.fiscot.org.

Hixon, M. A., Johnson, D. W., and Sogard, S. M. (2014). BOFFFFs: on the importance of conserving old-growth age structure in fishery populations. ICES Journal of Marine Science, 71:2171–2185. doi:10.1093/icesjms/fst200.

Hofbauer, J. and Sigmund, K. (1988). The theory of evolution and dynamical systems. London Mathematical Society Student Texts. Cambridge University Press, Cambridge UK.

ICES (2021). Tenth workshop on the development of quantitative assessment methodologies based on life-history traits, exploitation characteristics, and other relevant parameters for data-limited stocks (WKLIFE X). Technical Report ICES 2:98. 72 pp. http://doi.org/10.17895/ices.pub.5985.

Jennings, S., Pinnegar, J. K., Polunin, N. V. C., and Boon, T. W. (2001). Weak cross-species relationships between body size and trophic level belie powerful size-based trophic structuring in fish communities. Journal of Animal Ecology, 70:934–944.

Kleisner, K. and Pauly, D. (2011). The marine trophic index (MTI), the fishing in balance (FiB) index and the spatial expansion of fisheries. In Christensen, V., Lai, S., Palomares, M. L. D. Zeller, D., and Pauly, D., editors, The state of biodiversity and fisheries in regional seas, page 41–44. University of British Columbia, Fisheries Centre.

Kolding, J., Bundy, A., van Zwieten, P. A. M., and Plank, M. J. (2016). Fisheries, the inverted food pyramid. ICES Journal of Marine Science, 73:1697–1713. doi:10.1093/icesjms/fsv225.

Kvamsdal, S. F., Eide, A., Ekerhovd, N.-A., Enberg, K., Gudmundsdottir, A., Håkon Hoel, A., Mills, K. E., Mueter, F. J., Ravn-Jonsen, L., Sandal, L. K., Stiansen, J. E., and Vestergaard, N. (2016). Harvest control rules in modern fisheries management. Elementa: Science of the Anthropocene, 4:doi:10.12952/journal.elementa.000114.

Larkin, P. A. (1977). An epitaph for the concept of maximum sustained yield. Transactions of the American Fisheries Society, 106:1–11.

Law, R. and Plank, M. J. (2018). Balanced harvesting could reduce fisheries-induced evolution. Fish and Fisheries, 19:1078–1091. doi: 10.1111/faf.12313.

Law, R., Plank, M. J., and Kolding, J. (2012). On balanced exploitation of marine ecosystems: results from dynamic size spectra. ICES Journal of Marine Science, 69:602–614, doi:10.1093/icesjms/fss031.

Law, R., Plank, M. J., and Kolding, J. (2016). Balanced exploitation and coexistence of interacting, size-structured, fish species. Fish and Fisheries, 17:281–302. doi:10.1111/faf.12098.

Lewis, H. M. and Law, R. (2007). Effects of dynamics on ecological networks. Journal of Theoretical Biology, 247:64–76. doi:10.1016/j.jtbi.2007.02.006.

Link, J. S. and Watson, R. A. (2019). Global ecosystem overfishing: Clear delineation within real limits to production. Science Advances, 5:eaav0474. doi:10.1126/sciadv.aav0474.

Lubchenco, J. and Grorud-Colvert, K. (2015). Making waves: The science and politics of ocean protection. Science, 350:382–383. doi:10.1126/science.aad5443.

MacCall, A. D. (1980). The consequences of cannibalism in the stock-recruitment relationship of planktivorous pelagic fishes such as Engraulis. In Sharp, G. D., editor, Workshop on the effects of environmental variation on the survival of larval fishes, pages 201–220. Workshop Report 28: Intergovernmental Oceanographic Commission, UNESCO Paris.

Marañón, A. D., Cermeño, P., López-Sandoval, D. C., Rodríguez-Ramos, T., Sobrino, C., Huete-Ortega, M., Blanco, J. M., and Rodríguez, J. (2013). Unimodal size scaling of phytoplankton growth and the size dependence of nutrient uptake and use. Ecology Letters, 16:371–379. doi: 10.1111/ele.12052.

McKendrick, A. G. (1926). Applications of mathematics to medical problems. Proceedings of the Edinburgh Mathematical Society, 40:98–130.

Nilsen, I., Kolding, J., Hansen, C., and Howell, D. (2020). Exploring balanced harvesting by using an Atlantis ecosystem model for the Nordic and Barents Seas. Frontiers in Marine Science, 7:70. doi:10.3389/fmars.2020.00070.

Patterson, K. (1992). Fisheries for small pelagic species: an empirical approach to management targets. Reviews in Fish Biology and Fisheries, 2:321–338.

Pauly and Froese, R. (2020). MSY needs no epitaph—but it was abused. ICES Journal of Marine Science, page. doi:10.1093/icesjms/fsaa224.

Pauly, D., Christensen, V., Dalsgaard, J., Froese, R., and Torres Jr, F. (1998). Fishing down marine food webs. Science, 279:860–863. doi:10.1126/science.279.5352.860.

Pauly, D., Froese, R., and Holt, S. J. (2016). Balanced harvesting: The institutional incompatibilities. Marine Policy, 69:121–123. dx.doi.org/10.1016/j.marpol.2016.04.001.

Persson, L., Leeuwen, A. V., and Roos, A. M. D. (2014). The ecological foundation for ecosystem-based management of fisheries: mechanistic linkages between the individual-, population-, and community-level dynamics. ICES Journal of Marine Science, 71:2268–2280. doi:10.1093/icesjms/fst231.

Pikitch, E., Boersma, P. D., Boyd, I. L., Conover, D. O., Cury, P., Essington, T., Heppell, S. S., Houde, E. D., Mangel, M., Pauly, D., Plagányi, É., Sainsbury, K., and Steneck, R. S. (2012). Little fish, big impact: Managing a crucial link in ocean food webs. Technical report, Lenfest Ocean Program, Washington, DC.

Plank, M. J. (2018). How should fishing mortality be distributed under balanced harvesting? Fisheries Research, 207:171–174. doi.org/10.1016/j.fishres.2018.06.003.

Punt, A. E. (2010). Harvest control rules and fisheries management. In Grafton, R. Q., Hilborn, R., Squires, D., Tait, M., and Williams, M., editors, Handbook of Marine Fisheries Conservation and Management, pages 582–594. Oxford University Press, Inc, New York, NY.

Rehren, J. and Gascuel, D. (2020). Fishing without a trace? assessing the balanced harvest approach using EcoTroph. Frontiers in Marine Science, 7:510. doi:10.3389/fmars.2020.00510.

Ricker, W. E. and Foerster, R. E. (1948). Computation of fish production. Bulletin of the Bingham Oceanographic Collection, 11:173–211.

Rindorf, A., Dichmont, C. M., Levin, P. S., Mace, P., Pascoe, S., Prellezo, R., Punt, A. E., Reid, D. G., Stephenson, R., Ulrich, C., Vinther, M., and Clausen, L. W. (2017). Food for thought: pretty good multispecies yield. ICES Journal of Marine Science, 74:475–486. doi:10.1093/icesjms/fsw071.

Salomon, A. K., Gaichas, S. K., Jensen, O. P., Agostini, V. N., Sloan, N. A., Rice, J., McClanahan, T. R., Ruckelshaus, M. H., Levin, P. S., Dulvy, N. K., and Babcock, E. A. (2011). Bridging the divide between fisheries and marine conservation science. Bulletin of Marine Science, 87:251–274. doi:10.5343/bms.2010.1089.

San Martin, E., Irigoien, X., Harris, R. P., López-Urrutia, A., Zubkov, M. V., and Heywood, J. L. (2006). Variation in the transfer of energy in marine plankton along a productivity gradient in the Atlantic Ocean. Limnology and Oceanography, 51:2084–2091.

Scott, F., Blanchard, J. L., and Andersen, K. H. (2014). mizer: an R package for multispecies, trait-based and community size spectrum ecological modelling. Methods in Ecology and Evolution, 5:1121–1125. doi:10.1111/2041–210X.12256.

Sigmund, K. (1995). Darwin’s “circles of complexity”: assembling ecological communities. Complexity, 1:40–44.

Silvert, W. and Platt, T. (1978). Energy flux in the pelagic ecosystem: a time-dependent equation. Limnology and Oceanography, 23:813–816.

Sogard, S. M. (1997). Size-selective mortality in the juvenile stage of teleost fishes: a review. Bulletin of Marine Science, 60:1129–1157.

Soudijn, F. H., van Denderen, P. D., Heino, M., Dieckmann, U., and de Roos, A. M. (2021). Harvesting forage fish can prevent fishing-induced population collapses of large piscivorous fish. Proceedings of the National Academy of Sciences, 118:. doi.org/10.1073/pnas.1917079118.

Spence, M. A., Thorpe, R. B., Blackwell, P. G., Scott, F., Southwell, R., and Blanchard, J. L. (2021). Quantifying uncertainty and dynamical changes in multi-species fishing mortality rates, catches and biomass by combining state-space and size-based multi-species models. Fish and Fisheries, 22:667–681. doi:10.1111/faf.12543.

Svedäng, H. and Hornborg, S. (2017). Historic changes in length distributions of three Baltic cod (Gadus morhua) stocks: Evidence of growth retardation. Ecology and Evolution, 7:6089–6102. doi:10.1002/ece3.3173.

von Foerster, H. (1959). Some remarks on changing populations. In Stohlman, J. F., editor, The Kinetics of Cellular Proliferation, page 382–407. Grune and Stratton, New York.

Ware, D. M. (1978). Bioenergetics of pelagic fish: theoretical change in swimming speed and ration with body size. Journal of the Fisheries Research Board of Canada, 35:220–228.

Zhou, S., Kolding, J., Garcia, S. M., Plank, M. J., Bundy, A., Charles, A., Hansen, C., Heino, M., Howell, D., Jacobsen, N. S., ans JC Rice, D.G.R., and van Zwieten, P. A. M. (2019). Balanced harvest: concept, policies, evidence, and management implications. Reviews in Fish Biology and Fisheries, 29:711–733. doi.org/10.1007/s11160–019–09568–w.

Zhou, S. and Smith, A. D. M. (2017). Effect of fishing intensity and selectivity on trophic structure and fishery production. Marine Ecology Progress Series, 585:185–198. doi.org/10.3354/meps12402.

